# Mechanistic Insights into the Stimulation of the Histone H3K9 Methyltransferase Clr4 by Proximal H3K14 Ubiquitination

**DOI:** 10.1101/2024.09.28.615623

**Authors:** Yunxiang Du, Maoshen Sun, Zhengqing Li, Xiangwei Wu, Qian Qu, Huasong Ai, Lei Liu

## Abstract

H3K9 methylation is an evolutionarily conserved hallmark of heterochromatin and plays crucial roles in chromosome segregation, genome stability, and gene expression regulation. Clr4 is the sole histone methyltransferase responsible for catalyzing H3K9 methylation in *Schizosaccharomyces pombe*. Clr4 K455/K472 automethylation and histone H3K14 ubiquitination (H3K14Ub) are vital activators of the catalytic activity of Clr4, ensuring appropriate heterochromatin deposition and preventing deleterious gene silencing. While the mechanism by which automethylation activates Clr4 was recently elucidated, the mechanism of the significantly pronounced stimulatory effect of H3K14Ub on Clr4 remains unclear. Here we determined the crystal structures of Clr4 bound to ubiquitinated and unmodified H3 peptides at resolutions of 2.60 Å and 2.39 Å, respectively. Our structures reveal a synergistic mechanism underlying the stronger stimulatory effect by H3K14Ub compared to automethylation: site-specific ubiquitination constrained by the H3K14 linkage increases substrate affinity through multivalent interactions between ubiquitin and Clr4. Additionally, H3K14Ub facilitates the allosteric transition of Clr4 from an inactive apo conformation to a hyperactive “catalyzing state”, accompanied by conformational changes in the αC-SET-insertion (SI) region, complete release of the autoregulatory loop (ARL), and retraction of the β9/10 loop. Finally, we propose a structural model for the Clr4 catalytic-regulatory cycle, depicting varying levels of conformational regulation mediated by automethylation and ubiquitination. This work provides structural insights into the interplay between different histone modifications and their collective impact on epigenetic regulation.

## Introduction

Histone H3K9 methylation (H3K9me) is a central and conserved epigenetic hallmark of transcriptional silencing associated with heterochromatin formation^1–5^. H3K9 methylation deposition is a highly efficient process *in vivo*, as it induces a positive feedback loop involving chromodomain (CD)-mediated read-write process^6–8^ and crosstalk with RNAi-dependent co-transcriptional gene silencing^9^, leading to rapid H3K9me spreading. Multiple regulatory pathways are involved in controlling H3K9me generation, whose dysregulation is closely related to cancer, neurodegenerative, and viral diseases in humans^10–13^. Related pathways include H3K9 demethylation^14–18^, H3K9 acetylation occlusion and deacetylation^19–25^, and Argonaute (Ago1)-small RNA dependent^26–34^ or independent^25,35^ methyltransferase localization. These pathways precisely control H3K9me deposition and prevent aberrant heterochromatin deposition and deleterious gene silencing.

In addition to the above regulatory pathways, direct tuning of the histone methyltransferase (HMTase) activity that catalyzes H3K9me is indispensable for controlling H3K9me deposition. In *Schizosaccharomyces pombe* (*S. pombe*), which serves as a paradigmatic model organism for understanding the establishment and inheritance of heterochromatin, the sole H3K9 HMTase belonging to SUV39 superfamily, Clr4, undergoes automethylation at K455 and K472. This automethylation releases Clr4 from its intrinsic autoinhibitory state, which is necessary for appropriate heterochromatin establishment and epigenetic stability^36^. Besides being activated by automethylation, the HMTase activity of Clr4 is strongly promoted by H3K14Ub^37,38^, which is catalyzed by the Clr4-containing Clr4-Rik1-Cul4 methyltransferase complex (CLRC)^39–41^. The stimulatory effect of H3K14Ub on Clr4 is considerably more pronounced than that of the Clr4 automethylation^36,42^. Intriguingly, bioinformatics analysis suggested that Clr4 lacks a ubiquitin-binding domain, and experimental evidence indicates that it does not sense free ubiquitin^42^. The mechanism underlying the substantial stimulatory effect of H3K14Ub on Clr4 activity remains unknown.

Here, we determined the crystal structures of Clr4 bound to ubiquitinated and unmodified H3 peptides at resolutions of 2.60 Å and 2.39 Å, respectively. Combined with the results of biochemical experiments, our findings reveal the molecular mechanism underlying the synergistic effects of affinity and turnover enhancement that results in significant stimulation by H3K14Ub. Specifically, compared with the faint increase in substrate recruitment induced by Clr4 automethylation, H3K14Ub substantially enhances the affinity of Clr4 for substrate. The ubiquitin constrained by the isopeptide bond linkage at the H3K14 position forms multivalent interactions with the Clr4 catalytic domain, stabilizing substrate binding and rapidly promoting H3K9me2/3 deposition. Moreover, our findings indicate that the previously reported automethylated state of Clr4 requires additional allosteric changes to transition into the “catalyzing state”, which is characterized by conformational changes in the compact αC-SET-insertion (SI) region, complete release of the autoregulatory loop (ARL), and retraction of the β9/10 loop. Furthermore, H3K14 ubiquitination facilitates substrate turnover by promoting the transition of Clr4 to a hyperactive “catalyzing state”, primarily through interactions and conformational changes within the structural elements surrounding the β9/10 loop. Our work elucidates the continuous conformational changes occurring during the disinhibition of Clr4 via automethylation, substrate recruitment, and stimulation by H3K14Ub, thereby advancing our understanding of the regulation of H3K9 methylation.

## Results

### Functional interplay of Clr4 activation by H3K14Ub and automethylation

Our work began with biochemical reconstitution of the H3K14Ub-mediated stimulation of H3K9 methylation *in vitro* using full-length Clr4 (residues 1-490) or the Clr4 KMT domain (residues 192-490) with truncation of the N-terminal chromodomain (CD, residues 1-70) and the disordered hinge region (residues 71-191) **(Fig. 1b)**. H3K14 ubiquitinated and unmodified histone H3 N-terminal tail (residues 1-20) (hereafter referred to as Ub-H3t and H3t, respectively) were chemically synthesized as substrates **(Extended Data** Fig. 1a, **Supplementary** Fig. 1a**, 1b).** Histone methyltransferase activity assay of full-length Clr4 and the KMT domain on H3 tail substrates indicated that H3K14Ub significantly increased the production rate of S-adenosylhomocysteine (SAH) **(Extended Data** Fig. 1b**, 1e)**, validating the significant stimulatory effect of H3K14Ub on Clr4 methyltransferase activity^37,42^. Next, we explored the functional interplay between the activation of Clr4 methyltransferase activity by K455/K472 automethylation and H3K14Ub. The average methyl transfer rate per microMolar Clr4 calculated directly from the endpoint quantity of SAH showed that automethylation of Clr4 **(Supplementary** Fig. 2a**, 2b)** and automethylation-mimicking hyperactive mutations (K455A/K472A or K455W/K472W)^36^ activate the catalysis of both unmodified and ubiquitinated substrates **(Extended Data** Fig. 1b). In turn, the average methyl transfer rate of automethylated Clr4 can be further elevated by H3K14Ub for approximately 50 folds **(Extended Data** Fig. 1b). Collectively, the influences of the Clr4 automethylation and H3K14 ubiquitination are cumulative, collaborating to regulate Clr4 activity.

**Fig. 1.**
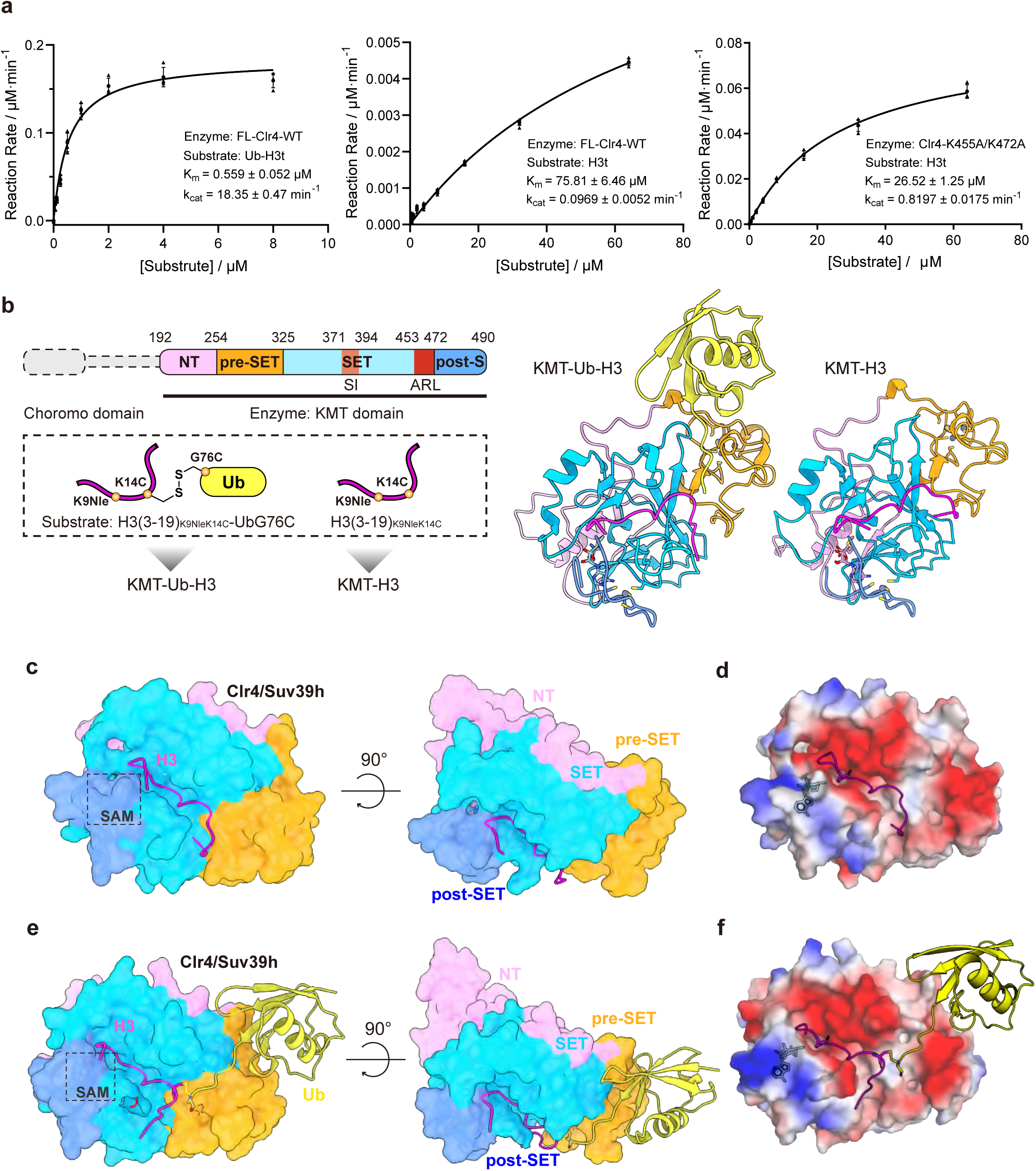
Biochemical reconstruction and global structures of complex KMT-Ub-H3 and KMT- H3. **a.** Michaelis-Menten curves of the full-length Clr4 methylation reaction with ubiquitinated (left) or unmodified (right) peptide as the substrate, and automethylation-mimicking Clr4 as the enzyme (n=4, error bars indicate SD). **b.** Construction of the complex KMT-Ub-H3 and KMT-H3, and their resulting crystal structures: KMT-H3 on the left and KMT-Ub-H3 on the right. Domains and subdomains in Clr4 (up), H3 (in magentas), and ubiquitin (in yellow) are distinguished by different colors, and the colors in this picture correspond to the colors of the surface in the following structure figures. **c. d.** Global structure of KMT-H3 complex and KMT-Ub-H3 complex, the surface of domain KMT is colored through different subdomains, histone H3 is shown in magentas and the Ub is shown in yellow. **e. f.** The surface electrostatic diagram of KMT in KMT-H3 and KMT-Ub-H3 structures, the H3 and Ub are still displayed in cartoon, and the SAM is represented by sticks.

Interestingly, the above measurements revealed that the activating effects of Clr4 automethylation and hyperactive mutations on methylation catalysis were significantly inferior to those of H3K14 ubiquitination. We conducted Michaelis-Menten kinetics experiments to analyze stimulatory effects quantitatively **(Fig. 1a)**. The Clr4 automethylation-mimicking mutation resulted in only a 2.9-fold increase in substrate affinity (*K*_m_ = 26.5 ± 1.2 μM vs. wild type *K*_m_ = 75.8 ± 6.5 μM) and an 8.5-fold enhancement of methylation turnover (*k*_cat_ = 0.820 ± 0.017 min^-1^ vs. wild type *k*_cat_ = 0.0969 ± 0.0052 min^-1^). In contrast, the H3K14Ub significantly enhanced both the substrate affinity by 135.6-fold (*K*_m_ = 0.559 ± 0.052 μM vs. unmodified *K*_m_ = 75.8 ± 6.5 μM) and turnover by 188.8-fold (*k*_cat_ = 18.3 ± 0.5 min^-1^ vs. unmodified *k*_cat_ = 0.0969 ± 0.0052 min^-1^) in H3K9 methylation, resulting in a 1061.5-fold increase in the specific constant *k*_cat_/ *K*_m_ (32.8 min^-1^·μM^-1^) compared with that of the Clr4 automethylation-mimicking mutation (0.0309 min^-1^·μM^-1^). These results indicated that the stimulatory effects of the reaction by H3K14Ub are more pronounced than that of Clr4 automethylation in terms of substrate affinity and intrinsic Clr4 activity.

### Crystal structures of Clr4 bound to H3K14Ub or unmodified H3 peptide

To elucidate the structural mechanism of the significant stimulatory effect of H3K14Ub on Clr4, we cocrystallized the catalytic KMT domain (composed of the NT, pre-SET, SET catalytic core, and post-SET subdomains) with the H3K14 ubiquitinated or unmodified H3 N-terminal tail peptide in the presence of S-adenosylmethionine (SAM). However, our initial attempts to crystallize the complex using a chemically synthesized homogeneous isopeptide-linked ubiquitinated H3K14 tail as a substrate were unsuccessful. After extensive optimization, including testing various covalent linkage strategies for ubiquitin and truncation of the H3 peptide, we succeeded in obtaining a crystal structure when we used a substrate mimic with an H3 N-terminal tail peptide (residues 3-19) and ubiquitin attached to K_C_14 via a disulfide bond **(Fig. 1b)**. This disulfide bond-mimicking strategy has been validated in previous ubiquitin-linked systems^43,44^. Specifically, K14 of H3 (3-19) was mutated to Cys to attach Ub_G76C_, and K9 was replaced with norleucine (Nle), which leads to SAM-dependent high affinity to Clr4 catalytic pocket and was widely used in MTase structural studies^45–47^ **(Extended Data** Fig. 1c**, Supplementary** Fig. 1c**, 1d)**. Compared to the native K14-ubiquitinated substrate, the disulfide bond-linked substrate exhibited comparable affinity in the isothermal titration calorimetry (ITC) test (*K*_D_ = 133 ± 13 μM for the disulfide-bond substrate versus *K*_D_ = 115 ± 18 μM for the isopeptide-bond substrate) **(Extended Data** Fig. 1d) and comparable stimulation efficiency **(Extended Data** Fig. 1e**, Supplementary** Fig. 1e). Finally, a 2.60 Å complex structure of the Clr4 KMT domain bound to H3(3-19)_K9Nle-K14C_-Ub_G76C_ **(Supplementary** Fig. 1f), and a 2.39 Å complex structure of the Clr4 KMT domain bound to H3(3-19)_K9Nle-K14C_ without Ub-modification **(Fig. 1b)** were resolved (hereafter abbreviated as the KMT-Ub-H3 and KMT-H3 structures, respectively) **(Supplementary** Fig. 3a-3f**, 4a-4h, Extended Data** **Table 1)**.

In these two complex structures, the H3-derived substrates and the SAM cofactor exhibit a 1:1:1 stoichiometry in their binding to the surface of the ovoid-shaped KMT domain. **(Fig. 1c, 1d)**. The KMT-Ub-H3 complex assembled into space group P2_1_2_1_2_1_ and KMT-H3 assembled into P22_1_2_1_. Each asymmetric unit of the two crystals contains two (P2_1_2_1_2_1_) copies and one (P22_1_2_1_) copy of the complex, respectively. For the crystal structure of the KMT-Ub-H3 complex, two copies the complex adopt highly similar conformations and interfaces **(Supplementary** Fig. 4a). In our structures, the KMT domain adopts a global conformation similar to that of the apo form (PDB: 6BOX)^36^ with an RMSD of ∼0.37 Å across 221 pairs of Cα. The H3 N-terminal tail peptide inserts into a histone-binding groove and straddles the SET subdomain of KMT, resulting in an interaction surface area of more than 2000 Å^2^. The section of the histone-binding groove near the catalytic active site is negatively charged since the acidic residue-rich SI region forms the outer “wall” of the groove and interacts with residues R8 to G13 of the H3 N-terminal tail **(Fig. 1e, 1f)**. The SAM cofactor is located on the larger side of the ovoid structure, embedded in the SAM pocket of the SET domain, with the post-SET subdomain covering above as a lid. Moreover, local allosteric changes are observed in certain structural elements such as the αC helices (residues 366-372), the SET-insertion (SI) region (residues 371-394), and the autoregulatory loop (ARL, residues 453-472).

Notably, in the KMT-Ub-H3 structure, the Ub moiety is clearly visible and docks with the KMT domain on the opposite side of the SAM-binding pocket, forming extensive interfaces, which are 1542 Å^2^ and 1559 Å^2^ in two NCS units, respectively. The C-terminal tail of Ub extends to the histone-binding groove along a neutral-hydrophobic canyon, sandwiched between the acidic pre-SET and SI region **(Fig. 1f)**.

### Conserved Clr4 residues in the histone-binding groove contribute to activity and selectivity

The N-terminal tail of histone H3 extensively interacts with the SET subdomain within the KMT domain. The majority of these contacts involve residues A7 to K_C_14 of H3, which are situated within a narrow groove formed by the parallel acidic SI region, the N-terminus of Clr4, and the initial segment (^451^YAGA^454^) of ARL **(Fig. 2a, 2b, Extended Data** Fig. 2a**, 2b)**. Within this groove, L382, D384, and D386 in the SI region form hydrogen bonds with the backbone of the H3 tail. The guanidine group of H3 residue R8 is further stabilized by hydrogen bonds with the side chains of D371 and D384 of Clr4. Additionally, the carbonyl oxygen of A367 and the hydroxy group of T395 interact with the guanidine group of R8, forming a quadrangular clamp structure that secures this residue (**Fig. 2a**). These interactions contribute to the substrate H3K9 specificity of Clr4, as the enzyme selectively recognizes an RK core motif within the histone tail. H3 K9_Nle_ is positioned within the catalytic pocket of Clr4, where its backbone along with the backbone of S10, forms main-chain hydrogen bonds with the SI region. The carboxy group of D386 further forms polar contacts with the hydroxy groups of H3 residues S10 and T11.

**Fig. 2.**
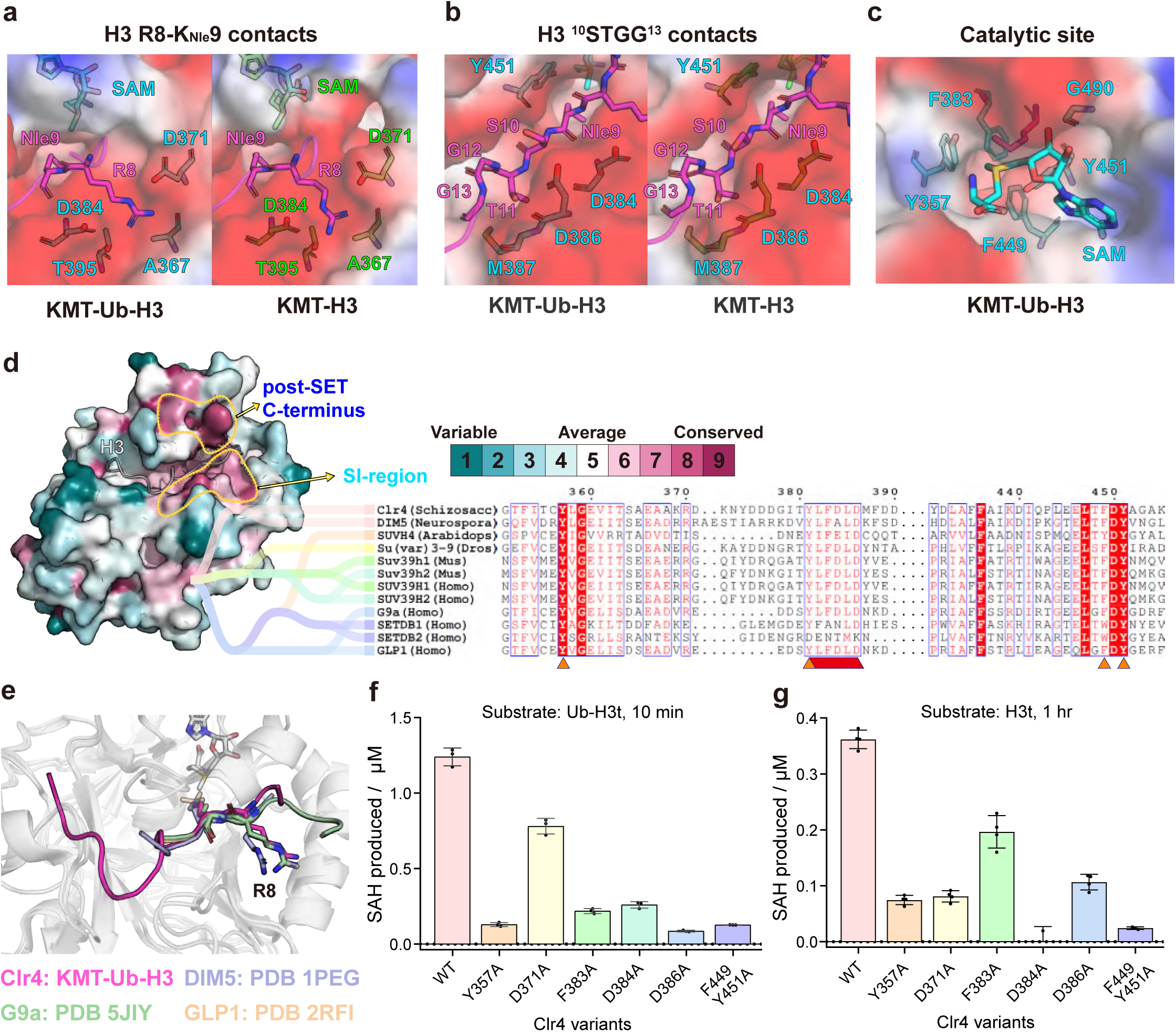
**Conserved interactions between Clr4 KMT and H3 N-terminal peptide**. **a.** Residues that form quadrangular clamp construct surrounding R8 and interact with the K_Nle_9 backbone. H3 is in magentas and residues of Clr4 in the KMT-Ub-H3 are represented by cyan sticks, while in KMT-H3 are green. **b.** Clr4 residues that contact with the H3 ^10^STGG^13^ in the KMT-Ub-H3 structure and KMT-H3 structure. **c.** Conformation of SAM and K_Nle_9 binding with the Clr4 catalytic pocket/site in KMT-Ub-H3 structure. **d.** Evolutionary conservation of residues in Clr4 between homologous proteins, conserved residues appear redder on the surface of KMT. The conserved SI-region and post-SET C-terminus are marked by the yellow circle. The right picture shows the evolutionary tree and sequence alignment between SUV39 family proteins, the conserved SI-region and catalytic pocket are marked by orange and red triangles. **e.** Comparison of H3 conformation among structures of Suv39 proteins binding with H3 N-terminal tail. H3 in our structure is in magentas, while the H3 binding with DIM5 (PDB: 1PEG) is in purple, binding with G9a (PDB: 5JIY) is in pale green, and binding with GLP1 (PDB: 2RFI) is in wheat. **f. g.** Chemiluminescence histone methyltransferase activity assay measuring SAH production during the WT Clr4 or Clr4 accommodating mutations in histone binding groove or catalytic site catalyzing substrate Ub-H3t and Ub methylation (n=3, error bars indicate SD), SDS-PAGE gels of the reactions are provided in the supplementary figures.

The side chain of histone H3 K9_Nle_ inserts into a hydrophobic tunnel and attacks the S-methyl group of SAM (**Fig. 2c**), which is consistent with the active site conformations of other H3K9- specific HMTase (**Extended Data** Fig. 2d). Four aromatic residues (Y357, F383, F449, and Y451) within the aromatic catalytic site surrounding the side chain of K9_Nle_ are highly conserved among various Clr4 homologs in eumycetes and higher living organisms, such as *Arabidopsis thaliana*, *Drosophila melanogaster*, *Mus musculus*, and *Homo sapiens* **(Fig. 2c, Extended Data** Fig. 2c). Besides the classic catalytic site, the acidic SI region is also conserved among H3K9-specific HTMases **(Fig. 2d)**. Compared with the substrate-binding structure of Clr4 homologs like DIM5 in *Neurospora crassa*^48^, G9a^49^, and GLP1^50^ in *Homo sapiens*, the conformation of H3 residues 7-13 in our structure is similar, and the SI region adopts a concerted recognition behavior when it is tightly interdigitated with residues 7-13 of the H3-tail peptide **(Fig. 2e)**. This observation suggests that the H3K9-specific HTMases preserve H3K9 recognition patterns and selectivity across various species throughout evolution.

To validate the interfaces between Clr4 and H3, we introduced single or double alanine mutations at key interface residues of Clr4 and tested the catalytic behavior of these mutants using unmodified or ubiquitinated H3 N-terminal tail (1-20) (H3t and Ub-H3t). We found that alanine mutations of either SI region residues (D371, D384, and D386), whose side chains are involved in hydrogen bond formation, or residues at the catalytic site (Y357, F383, F449, and Y451) weakened the methylation activity, regardless of whether the substrate was unmodified H3 or ubiquitinated H3 **(Fig. 2f, 2g, Supplementary** Fig. 5a**, 5b)**. Therefore, our structural and biochemical results indicate that the highly conserved interactions between the Clr4 histone-binding groove and H3 (residues 7- 14) are critical for selective substrate binding and methyltransferase activity.

### Clr4-Ub interfaces are crucial for the stimulation

We next focused on the structure of the KMT-Ub-H3 complex, in which the Ub moiety resembles a balloon tied to a string **(Fig. 3a, Supplementary** Fig. 4b). The interfaces between Ub and Clr4 can be categorized into three distinct regions **(Fig. 3a)**: First, the canonical Ub interacting spots, including residues L8, I44, V70 and the aliphatic portion of the K6 side chain, create a hydrophobic pocket that accommodates F256 on Clr4 loop 2. Clr4 residue F256 is further stabilized by π-π interaction with Ub residue H68 **(Fig. 3b, Extended Data** Fig. 3a). Second, R42 and Q49 of Ub are inserted into an acidic pocket of the pre-SET domain. R42 forms salt bridge interactions with Clr4 D281 and forms a hydrogen bond with S258, whereas Q49 contacts with Clr4 D280 **(Fig. 3c, Extended Data** Fig. 3b). The third interface is at the C-terminal tail of Ub (residues 70-76), whose main chain along with the side chains of V70 and L73, lies in the neutral-hydrophobic canyon, which starts from F256 of Clr4 and extends to the histone-binding groove. Meanwhile, the side chain of R72 is fastened to the acidic pre-SET base through two salt bridge interactions with D280 and D281 **(Fig. 3d, Extended Data** Fig. 3c).

**Fig. 3.**
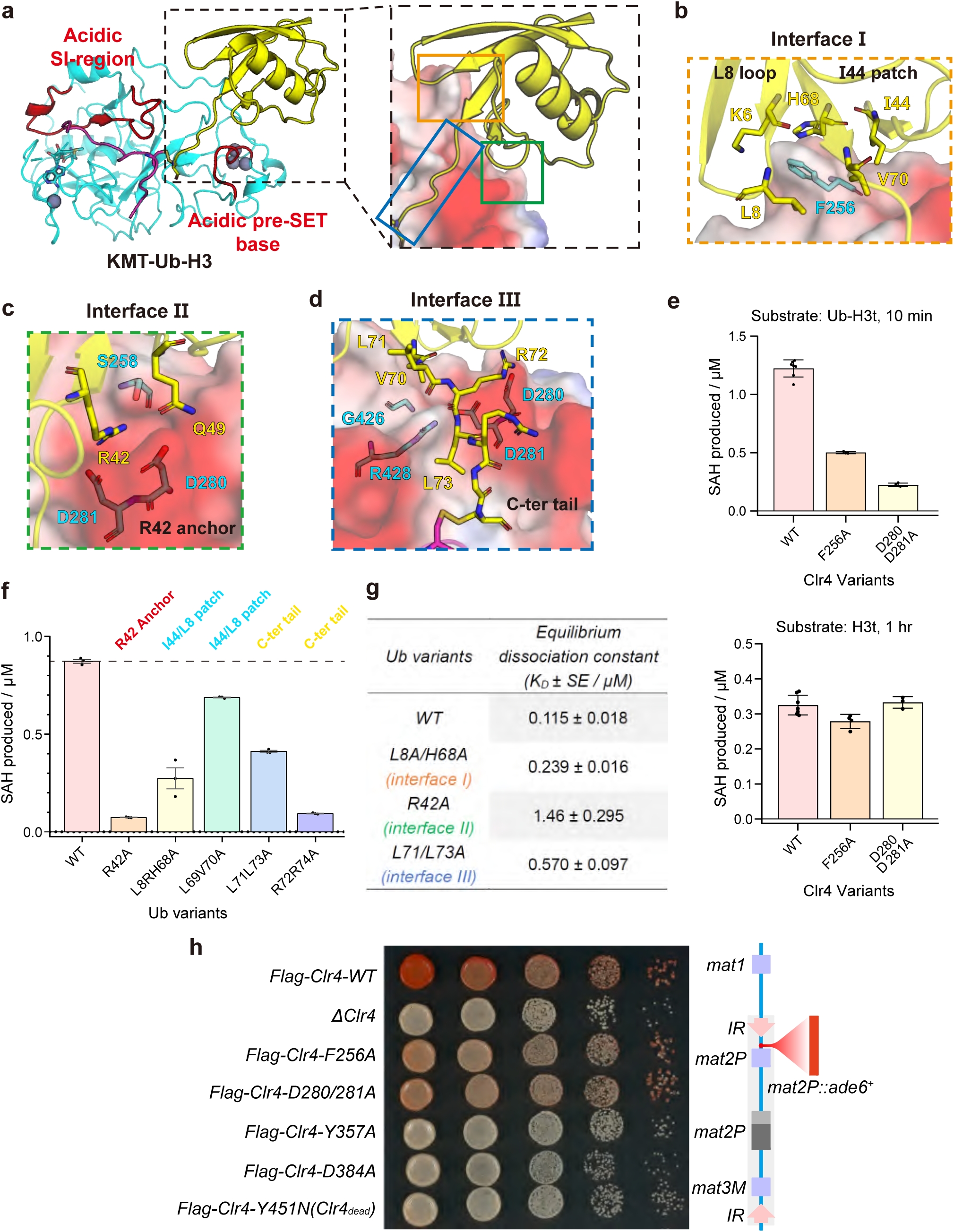
Multivalent binding between ubiquitin and Clr4 KMT. **a.** Multivalent binding behavior between ubiquitin and Clr4 KMT, ubiquitin is shown in yellow, Clr4 is shown in cyan while the key acidic regions are highlighted in red. **b. c. d.** Close-up views of three interfaces (I, II, and III) between Ub and KMT. **e.** Histone methyltransferase activity assay of WT Clr4 or Clr4 accommodating mutations in ubiquitin interfaces working on substrate Ub-H3t (up) and Ub (down). **f.** Histone methyltransferase activity assay of WT Clr4 working on Ub-mutated Ub-H3t. **g.** *K*_D_ value measured by ITC experiment performed with WT Clr4 KMT and Ub interface-mutated Ub-H3t. h. silencing assays of *mat2P::ade6^+^* on low-adenine medium (Low ade) to assess *ade6^+^* silencing in the indicated genotypes involving Clr4 and Clr4 mutants.

The importance of the Clr4-Ub interface was assessed by evaluating the impact of Clr4 mutations. The Clr4 F256A mutation, which diminishes hydrophobic interactions, reduced the histone methyltransferase activity of Ub-H3t by 60% (**Fig. 3e** **up, Supplementary** Fig. 5c).

Mutations in the pre-SET acidic pocket (D280A/D281A) strongly impaired histone methyltransferase activity by 82%. In contrast, these Clr4 mutants (F256A, and D280A/D281A) did not affect H3K9 methylation activity on unmodified H3t **(Fig. 3e** **bottom, Supplementary** Fig. 5c). We also conducted thermoshift experiments and revealed that the ΔTm of these mutants significantly decreased when binding to the ubiquitinated H3 tail, indicating the importance of these interfaces for ubiquitin binding and thermodynamic stabilization of Clr4 **(Extended Data** Fig. 3e). Furthermore, Clr4 methylation activity was noticeably decreased on the Ub-H3t substrate bearing mutations in the Ub R42 anchor (R42A), I44/L8 patch (L8R/H68A, L69A/V70A), and C-terminal tail (L71A/L73A, R72A/R74A). Mutation of R42 led to the most significant decrease in the ability to be methylated (92%), while that of the other four mutants was reduced by 22∼89% **(Fig. 3f, Supplementary** Fig. 5d**, 6a-e)**. Further measurement of the binding affinity between the above Ub-H3t variants and Clr4 by ITC revealed dissociation constant *K*_D_ values ranging from 0.239 to 1.46 μM, which are much lower than that of the wild-type substrate (0.115 ± 0.018 μM) **(Fig. 3g, Extended Data** Fig. 3d). Collectively, these findings indicate that the interfaces between Clr4 and H3K14 ubiquitin are crucial for substrate recognition, enzyme stabilization, and methylation stimulation.

### Validation of the key roles of the Clr4-H3 and Clr4-Ub interfaces *in vivo*

To validate the pivotal roles of the Clr4-H3 and Clr4-Ub interfaces identified in our crystal structure in the context of Clr4 function *in vivo*, we constructed *S. pombe* cells that express endogenous FLAG-tagged Clr4 mutant proteins with single or double substitutions at the Clr4-H3 (Y357A, and D384A) and Clr4-Ub interfaces (F256A, and D280A/D281A) **(Supplementary** Fig. 6f). Clr4-mediated heterochromatin deposition and gene silencing were examined using an *ade6^+^* reporter transgene inserted within the heterochromatic *mat* locus. The propagation of surrounding heterochromatin leads to the silencing of *ade6^+^*, resulting in red colonies on low-adenine plates. Our experiments showed that disruption of the catalytic pocket (Y357A) and histone-binding groove (D384A) completely abolished the heterochromatin deposition, yielding colonies as white as those observed with the previously reported Clr4 dead mutant (Y451N)^42^ **(Fig. 3h)**. This outcome underscores the essential roles of both the histone-binding groove and Clr4 methylation activity in Clr4-mediated heterochromatin deposition. Additionally, mutations at the Clr4-Ub interfaces resulted in a significant decrease in silencing, as evidenced by light pink to white colonies **(Fig. 3h)**. These findings indicate that the Clr4-Ub interfaces are important for the regulation of heterochromatin deposition, while the disruption of Ub interactions still allows minimal heterochromatin formation.

### Essentiality of the H3K14-specific junction for ubiquitin-mediated stimulation

Notably, the interaction between Clr4 KMT and free ubiquitin was undetectable in the ITC experiment **(Extended Data** Fig. 3f), which is consistent with previous findings that Clr4 does not sense free ubiquitin to facilitate H3K9 methylation^42^ **(Fig. 4a, Supplementary** Fig. 7a). These findings suggests that ubiquitin’s stimulatory effect of ubiquitin is contingent on its linkage to H3K14. We therefore hypothesized that the binding of ubiquitin to KMT is constrained by the H3K14 junction, which in turn enhances binding affinity by stabilizing the enzyme-substrate complex. In alignment with this hypothesis, the presence of free ubiquitin did not influence the change in the melting temperature (ΔTm) of the complex in the thermal shift assay **(Fig. 4b, Extended Data** Fig. 4a**, Supplementary** Fig. 7c**, 7d)**. Furthermore, extending the C-terminal tail of ubiquitin with an AEEA ([2-(2-aminoethoxy) ethoxy] acetic acid) linker **(Extended Data** Fig. 4c) led to an approximately 25% reduction in methyltransferase activity with the Ub-AEEA-H3t substrate **(Fig. 4a)** and a 1.12 °C decrease in ΔTm compared with that of the native Ub-H3t substrate **(Fig. 4c, Extended Data** Fig. 4b**, Supplementary** Fig. 7b). This result suggested that a less constrained ubiquitin motif resulted in reduced thermodynamic stabilization and consequently weaker activation of Clr4.

**Fig. 4.**
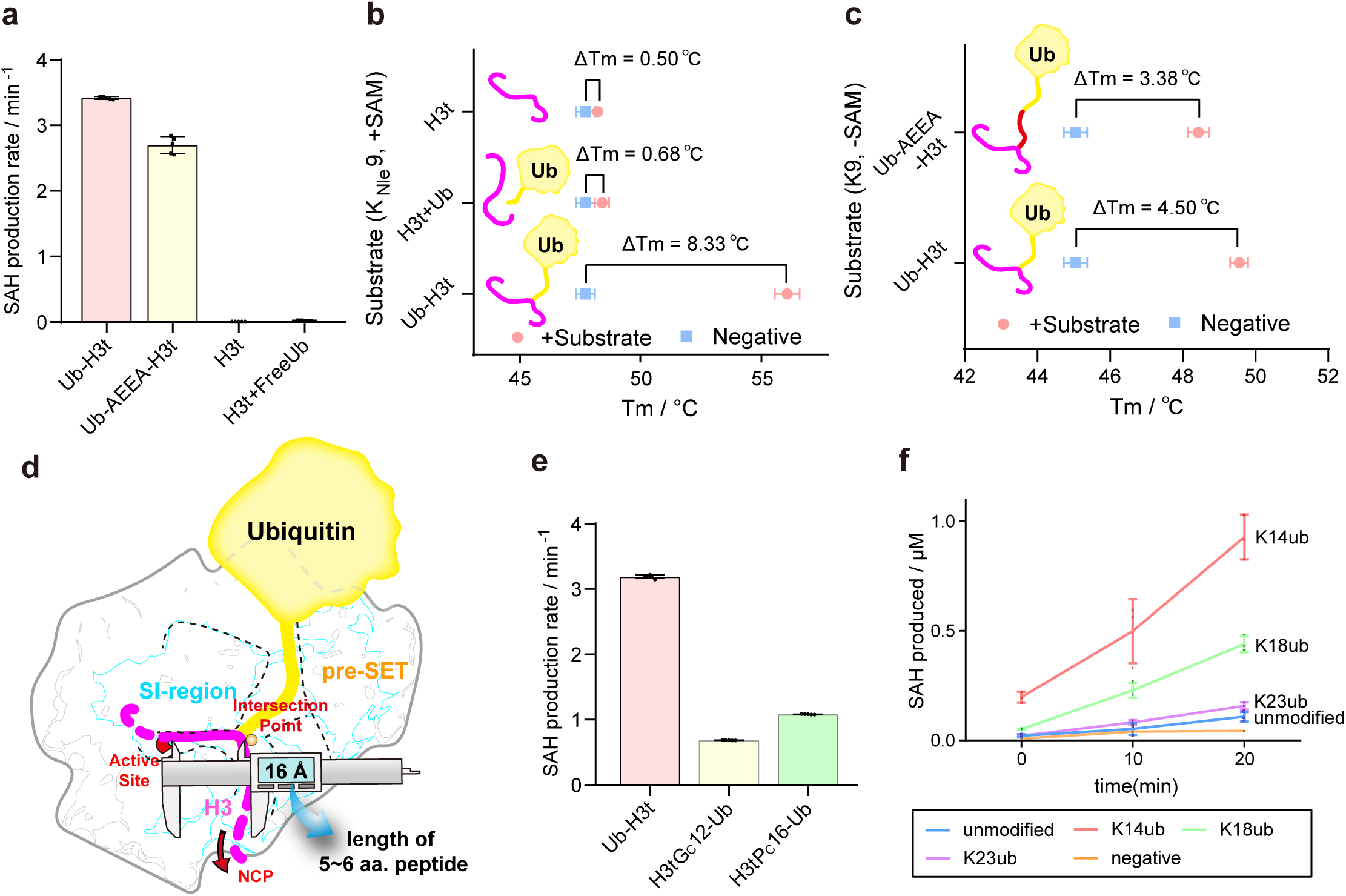
Biochemical experiment about H3-Ub junction. **a.** Histone methyltransferase activity assay of WT Clr4 working on the substrates with or without ubiquitination and AEEA-linked ubiquitination (n=5, error bars indicate SD). **b.** Thermo-shift assay working on Clr4 KMT binding with Ub-H3t(K_Nle_9), H3t(K_Nle_9), H3t(K_Nle_9) with free Ub. The melting temperature was determined by the first derivative of the barycentric mean (BCM) of fluorescence intensity with respect to temperature (dBCM/dT) (n=3, error bars indicate SD). **c.** Thermo-shift assay performed with Clr4 KMT and Ub-H3t, Ub-AEEA-H3t, respectively (n=3, error bars indicate SD). **d. S**cheme of “caliper construct” functioning in the recognition of K14-ubiquitinated H3 N-terminal tail. The catalytic site is highlighted by a red circle and the intersection point of the histone-binding groove and neutral- hydrophobic canyon is labeled by an orange dot. The caliper shows the approximate distance between the catalytic site and the intersection point. **e.** Histone methyltransferase activity assay of WT Clr4 working on the substrates that ubiquitin linked to residue 13 or residue 16 of H3 N-terminal tail (n=5, error bars indicate SD). **f.** Histone methyltransferase activity assay of WT Clr4 catalyzing NCP substrate with H3K14/K18/K23Ub or without modification (n=3, error bars indicate SD).

In our KMT-Ub-H3 complex structure, the distance from the catalytic site to the point where the histone-binding groove intersects with the neutral-hydrophobic canyon is approximately 16 Å. This distance is slightly shorter than that of an extended hexapeptide (∼19 Å, corresponding to residues ^9^KSTGGK^14^), as the sequential Cα−Cα distance is consistently restricted to ∼3.8 Å **(Extended Data** Fig. 4e). Notably, the K14 junction is located at this intersection, allowing the histone-binding groove to effectively accommodate the extended configuration of H3 residues K9 to K14 while enabling the C-terminal tail of Ub to reside precisely within the neutral-hydrophobic canyon **(Fig. 4d)**. This recognition mechanism implies that the stimulatory effect is highly sensitive to the position of the ubiquitination site. Attempts to fine-tune the ubiquitination site, either upstream or downstream of H3K14, led to impaired methylation activity with the Ub-modified H3 N-terminal tail **(Fig. 4e, Supplementary** Fig. 8a), despite only shifting the junction by 2 amino acids forward or backward (H3tG_C_12-Ub, H3tP_C_16-Ub, **Supplementary** Fig. 8c**, 8d**). This finding highlights the low tolerance for alterations in the substrate ubiquitination site.

Furthermore, we investigated biologically relevant ubiquitinated H3 substrates, including K18 and K23-ubiquitinated nucleosomes involved in DNMT1 recruitment and activation in the context of hemimethylated DNA inheritance^51–53^ (NCP_H3K14Ub_, NCP_H3K18Ub_, and NCP_H3K23Ub_, **Supplementary** Fig. 8b**, 8e-8g**). Histone methyltransferase assay revealed that the methylation rates of unmodified NCP, NCP_H3K18Ub_, and NCP_H3K23Ub_ were much lower than that of NCP_H3K14Ub_ **(Fig. 4f)**. The electrophoretic mobility shift assay (EMSA) results indicated that almost all NCP_H3K14Ub_ was bound to Clr4 at a Clr4 concentration of 0.5 μM, whereas a portion of NCP_H3K18Ub_, along with unmodified NCP, persisted as free NCP until the concentration of Clr4 was increased to 2 μM **(Extended Data** Fig. 4d). These results imply that Clr4 is selectively recruited to H3K14-ubiquitinated nucleosomes.

### Allosteric effects during substrate turnover and its stimulation by H3K14Ub

Next, we compared the structures of Clr4 in the KMT-H3, KMT-Ub-H3, and apo form, to analyze the conformational changes mediated by histone H3 and Ub respectively, which underlie the pronounced difference in Clr4 substrate turnover (*k*_cat_) observed during catalysis of unmodified and H3K14Ub-modified substrates.

We first superimposed our KMT-H3 complex structure with the previously reported Clr4 KMT apo conformation structure (PDB: 6BOX) to identify key conformational changes required for substrate engagement. Structural rearrangements of Clr4 were observed in four main regions: the ARL (residues 453-472), the helix αC-SI region (residues 365-395), the Clr4 C-terminus (residues 486-490), and the β9/10 loop (residues 423-431) **(Extended Data** Fig. 5a**, 5b)**. Once H3 binds, the ARL, which previously occluded the active site^36^, is released to expose the active site and accommodate the binding of the H3 N-terminal tail (**Fig. 5b**). The Clr4 C-terminus undergoes a ∼75° rotation with G486 as the vertex, positioning itself toward the substrate **(Fig. 5b)**. Helix αC (residues 364-472) is rotated approximately 9° toward the substrate on the axis of T363, generating a hydrogen bond between Clr4 D371 and H3 R8 **(Fig. 5a)**. The β-hairpin loop within the SI region (residues 381-394) shifts toward the histone-binding groove by 1.62 Å backbone-RMSD to interact with H3 **(Fig. 5c)**. Meanwhile, the β9/10 loop, located at the center of the neutral-hydrophobic canyon and in close proximity to the β-hairpin loop (residues 385-394) of SI region, undergoes retraction in the “catalyzing state”, which facilitates the formation of a new salt bridge interaction between Clr4 residues R428 and D390 on the β-hairpin loop in SI region **(Fig. 5c)**. These observations suggest that specific conformational changes within structural elements of Clr4 are required for the positioning and alignment of H3 for methylation.

**Fig. 5.**
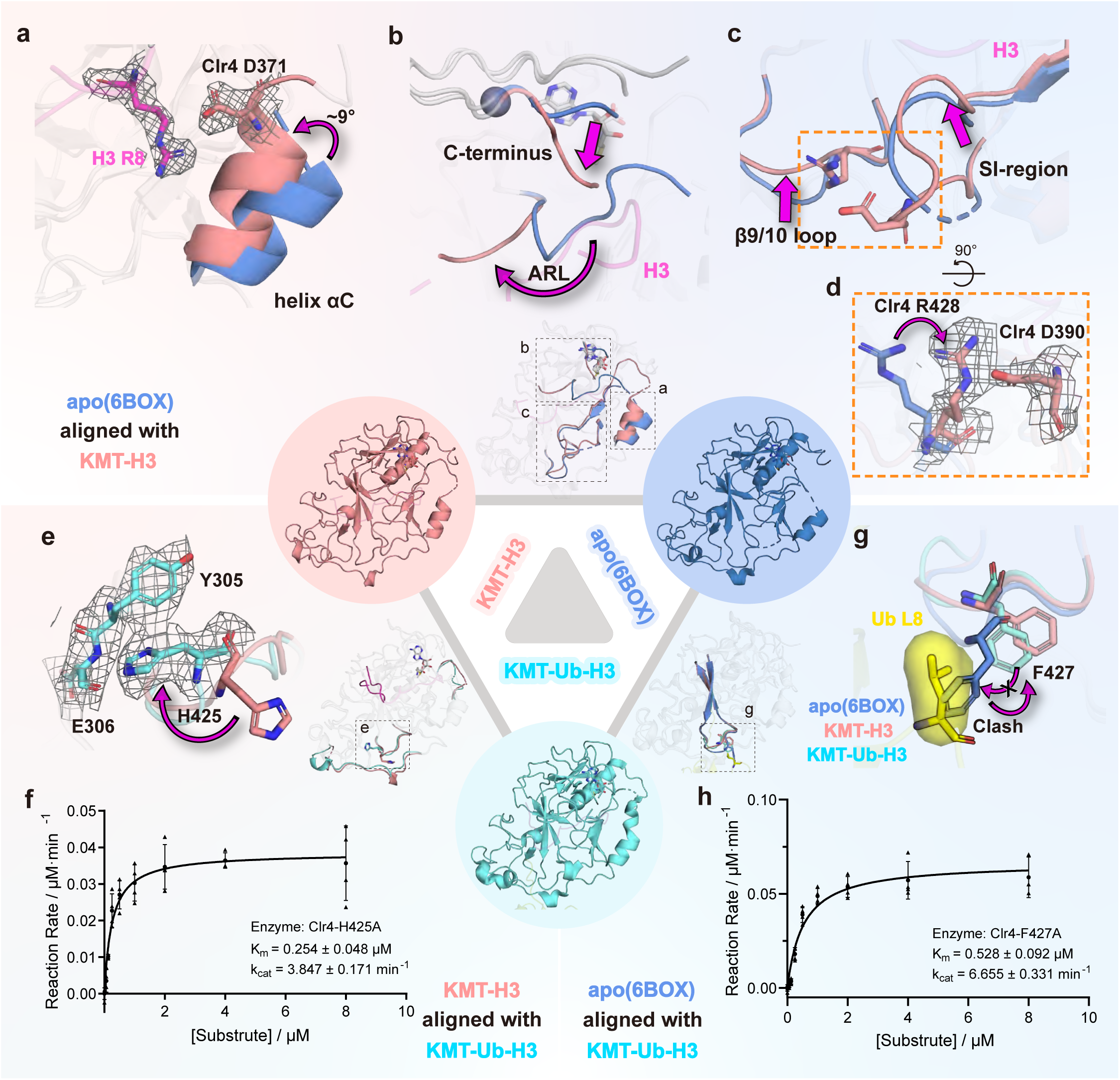
**Allostery effects during substrate turnover and stimulation by ubiquitin.** **a.** Conformation changes of helix αC between apo structure and our KMT-H3 structure. Two structures are superposed by KMT alignment. Structural elements in the apo structure are colored in blue and those in the KMT-H3 structure are colored in pink. The directions of allostery are highlighted by magenta arrows. Clr4 D371 and H3 R8 forming new interactions are shown as sticks in map density. **b.** Conformation changes of ARL and Clr4 C-terminus between apo structure and our KMT-H3 structure. **c, d.** Conformation changes of β9/10 loop and SI region between apo structure and our KMT-H3 structure. Clr4 D390 and R428 forming new interactions are shown as sticks in map density in **d)**. **e.** Flipping of Clr4 H425 once the H3K14 is ubiquitinated. Key residues forming new contacts are represented by sticks in map density. KMT-H3 structure and our KMT-Ub-H3 structure are superposed by KMT alignment, The KMT-H3 structure is colored in pink, and that in the KMT-Ub-H3 structure is colored in cyan. **f. h.** Michaelis-Menten curves of the full-length Clr4 accommodate **f)** H425A and **h)** F427A mutations catalyzing ubiquitinated peptide methylation (n=3, error bars indicate SD). **g.** Clash between Ub L8 and Clr4 F427 when superposing apo form Clr4 into the KMT-Ub-H3 structure. Ubiquitin is colored in yellow and L8 is represented by yellow sticks and surface.

To further understand how H3K14Ub drives the establishment of a hyperactive state of Clr4, we compared the KMT-H3 complex structure with the KMT-Ub-H3 complex structure to investigate the conformational change induced by H3K14 ubiquitination, which revealed structural changes in Clr4 at the β9/10 loop (residues 423-431), the beginning of the pre-SET subdomain (residues 254-268), and the onset of the SI region (residues 372-378), as well as structural changes at H3 residues 15-19 **(Extended Data** Fig. 5c**, 5d)**. Specifically, H425 in the β9/10 loop, which is well resolved in both the KMT-H3 and KMT-Ub-H3 structures, undergoes a 180° flip in the KMT-Ub-H3 structure. This flip allows it to engage in a π-π interaction with the aryl group of pre-SET residue Y305 and form a polar interaction with the backbone carbonyl oxygen of E306 **(Fig. 5e)**. This interaction may stabilize the retracted conformation of the β9/10 loop in the ubiquitin-induced hyperactive state. Further experiments involving the Clr4 H425A mutation demonstrated a significant reduction in *k*_cat_ for the Ub-H3t substrate, with an approximately 5-fold decrease (*k*_cat_ = 3.85 ± 0.171 min^-1^ vs. *k*_cat_ = 18.3 ± 0.5 min^-1^ for WT). Importantly, this mutation did not diminish the substrate recruitment (*K*_m_ = 0.254 ± 0.048 μM vs. *K*_m_ = 0.559 ± 0.052 μM for WT) or affect methylation activity with unmodified H3t (**Fig. 5f, Extended Data** Fig. 5e). These results suggest that the allostery triggered by H3K14Ub and the Clr4 β9/10 loop, along with the induced interactions between this loop and the pre-SET region, play essential roles in enhancing enzymatic turnover.

To identify the requisite conformational alterations that Clr4 undergoes upon binding to the H3K14Ub substrate, we compared the inactive Clr4 KMT apo structure (PDB: 6BOX) with the hyperactive KMT-Ub-H3 structure. This comparison revealed a steric clash between the Ub L8 loop and apo-form Clr4 residue F427 (**Fig. 5g**). In both the H3-KMT and H3-Ub-KMT structures, F427 is in a retracted position, forming a nascent hydrophobic pocket that accommodates the Ub L8 loop. This arrangement, in turn, restricts the β9/10 loop to the retracted conformation, which may be essential for the hyperactive state of the H3-Ub-KMT structure (**Fig. 5g**). Further analysis revealed that the Clr4 F427A mutation, which reduces the spatial size of the side chain, results in an approximately one-third reduction in *k*_cat_ (*k*_cat_ = 6.66 ± 0.33 min^-1^ vs. *k*_cat_ = 18.3 ± 0.5 min^-1^ of WT), while it does not significantly affect *K*_m_ (*K*_m_ = 0.528 ± 0.092 μM vs. *K*_m_ = 0.559 ± 0.052 μM of WT) or basal methylation activity with unmodified H3t (**Fig. 5h**, **Extended Data** Fig. 5e). These findings support the hypothesis that the newly formed interaction between Clr4 F427 and Ub is critical for driving H3K14Ub-mediated activation of enzymatic turnover.

## Discussion

Our study elucidates the molecular mechanism by which H3K14Ub significantly enhances the catalytic activity of Clr4 in the context of H3K9 methylation, a process central to heterochromatin formation^1,4,5^. Our structural and biochemical analyses revealed a synergistic mechanism involving both increased substrate affinity and allosteric activation **(Fig. 6b)**. First, H3K14Ub dramatically increases Clr4’s affinity for histone substrate through multivalent hydrophobic patches and salt bridge interactions between the Clr4 pre-SET subdomain and ubiquitin. The increase in enzyme-substrate binding is highly specific to H3K14 ubiquitination, highlighting the critical role of this precise modification site. Second, H3K14Ub triggers a conformational shift in Clr4, transitioning it to a hyperactive catalyzing state. This state is characterized by a compacted αC helix-SI region, complete release of the autoregulatory loop, and stable retraction of the β9/10 loop, which contribute to increased substrate turnover. In contrast, Clr4 K455/K472 automethylation, while contributing to Clr4 activity, demonstrates a less pronounced effect in comparison to H3K14Ub. Collectively, H3K14Ub acts as a potent dual-function regulator of Clr4, significantly boosting its activity through enhanced substrate binding and allosteric modulation, thereby playing a crucial role in the efficient establishment of heterochromatin.

**Fig. 6.**
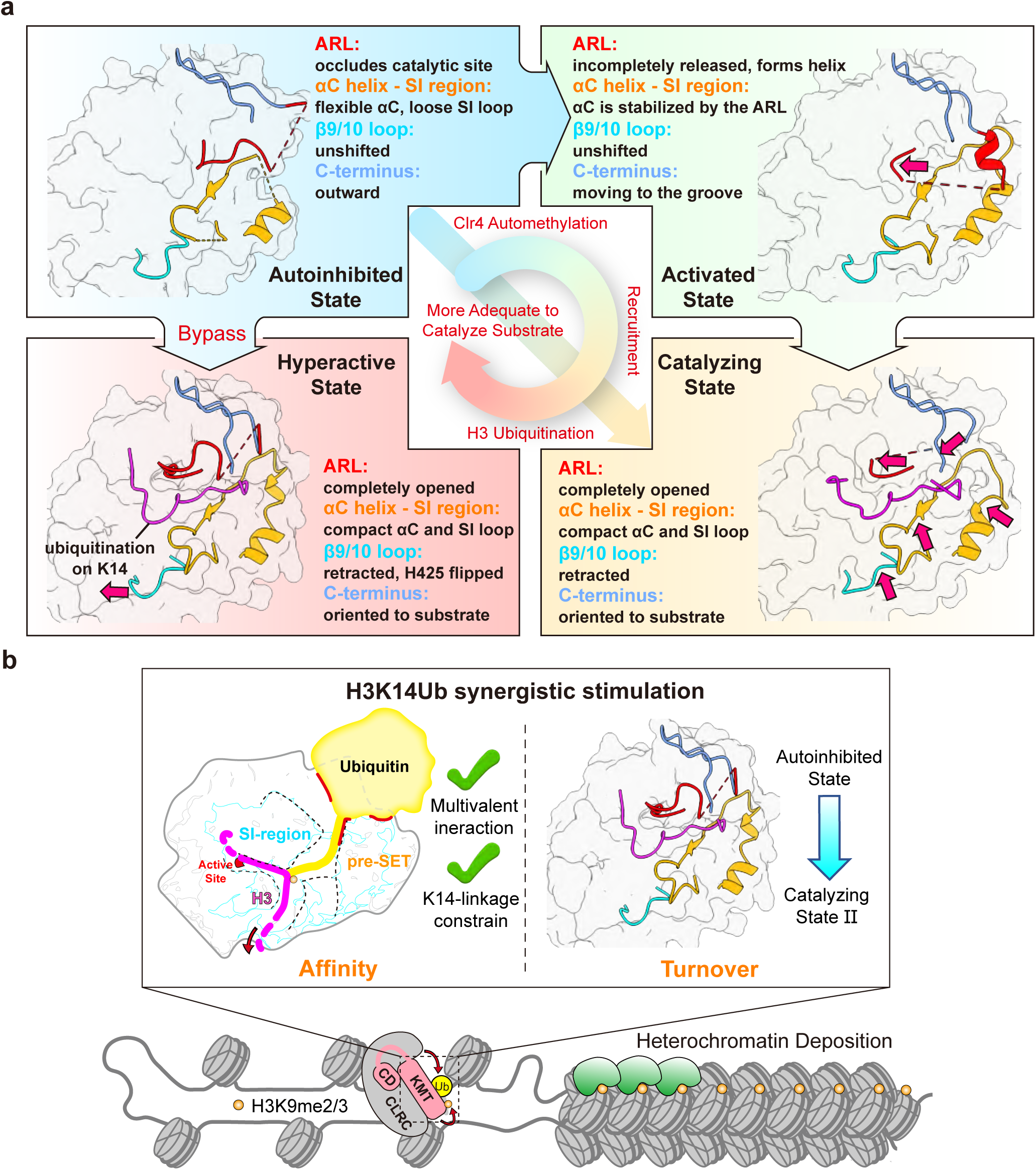
Multilayered regulation of Clr4 KMT catalyzation. **a.** Overview of conformation changes in the regulatory and catalyzation of Clr4. The element colors in cartoon schematic diagrams of structures correspond to the font fill colors of their names. **b.** Scheme of the synergetic mechanism of Clr4 stimulation involving both increased substrate affinity and allosteric activation by H3K14Ub.

By aligning our structures of “catalyzing state” and ubiquitin-induced “hyperactive state” of Clr4 with the previously resolved^36^ structures of Clr4 before (“autoinhibited state”, PDB: 6BOX) and after (“activated state”, PDB: 6BP4) automethylation, we propose the perspective of different levels of conformational regulation mediated by these two pathways **(Fig. 6a)**. Compared with the apo “autoinhibited state”, Clr4 in the “activated state” undergoes the relief of the ARL from the histone-binding groove and stabilization of the helix αC-SI region mediated by the ARL nascent C-terminal helix (residues 468-473) **(Extended Data** Fig. 5a). However, the helix αC-SI region and β9/10 loop remain in unaltered original conformations in the “activated state”, which is different from the “catalyzing state” **(Extended Data** Fig. 5a**, 5b)**. More importantly, the ARL in the “activated state” structure is supposed to be further opened when Clr4 is transferred to the “catalyzing state”. Because the ARL N-terminus (residues 451-455) in the “activated state” structure remains a steric clash with substrate H3 **(sFig. 10f)**, and the ARL C-terminal helix R469, I470 would impede the conformational shift of D371, K372, and the formation of D371-R8 hydrogen bonds **(Extended Data** Fig. 5c). Briefly, automethylation partially alleviates the clashes but cannot completely remove them. However, H3K14Ub induces a “hyperactive state”, involving the retracted β9/10 loop by the H425-pre-SET Y305/E306 and F427-Ub L8 interactions, as well as the shifted SI region contacting with β9/10 loop. Compared with automethylation, H3K14Ub adopts a distinct and likely more adequate mechanism to allosterically promote substrate turnover, contributing to the pronounced stimulation of H3K9 methylation.

The identification of Clr4 ubiquitin-binding interfaces provides valuable insights into the regulation of ubiquitination during gene silencing and heterochromatin formation. We found that mutation of F256 and D280/D281 of Clr4 significantly reduces the stimulatory effect of H3K14Ub on Clr4 activity without affecting its intrinsic H3K9 methylation activity **(Fig. 3e)**. The combined mutations of these amino acids present an ideal strategy for uncoupling the crosstalk between H3K14Ub and H3K9me, enabling the elucidation of the functional implications of this crosstalk for the downstream pathways associated with H3K9 methylation. For example, to investigate the function of H3K4 demethylation affected by H3K14ac, a recent study resolved the structure of the demethylase complex LSD1-CoREST recognizing H3K14ac, engineered a protein mutant that can uncouple this crosstalk, and clarified the role of this crosstalk in cellular adhesion and myeloid leukocyte activation^54^. The Clr4 mutations identified here are promising functional mutants for future biological studies.

Clr4-mediated H3K9 methylation creates binding sites for chromodomain-containing proteins such as Swi6^17,55,56^, facilitating the formation of a repressive chromatin structure. Once a threshold level of H3K9 methylation is achieved, Clr4 may rapidly diffuse across chromatin through a “read-write” mechanism mediated by the chromodomain to establish heterochromatin and gene silencing. Our biochemical and structural analysis suggested that H3K14 ubiquitination may play a critical initiator role in the early establishment of H3K9 methylation in fission yeast. Specifically, 1) the CLRC ubiquitin ligase complex is recruited by Ago1/RNAi-related complexes such as RNA-induced transcriptional silencing (RITS) complex, facilitating histone H3K14 ubiquitination at specific loci while assisting in the recruitment of Clr4^30–34^. 2) H3K14Ub can directly stimulate Clr4 activity irrespective of whether Clr4 is in an intrinsic autoinhibited state or not. 3) H3K14Ub enables Clr4 to efficiently override the rate-limiting step from dimethylation to trimethylation^8^, which produces the suitable substrate for the Clr4 chromodomain, promoting the rapid accumulation of H3K9me3 and enhancing subsequent H3K9me diffusion **(Extended Data** Fig. 6d). 4) Ubiquitination is a relatively complicated and consumable process involving ATP, E1 activating enzymes, E2 conjugating enzymes, E3 ligases, and ubiquitin. It seems difficult for ubiquitin to function as an extensive factor in the rapid H3K9me diffusion process. Conversely, it is likely to be more suitable as the “gating” for H3K9 deposition. The integration of these regulatory pathways illustrates a sophisticated mechanism ensuring precise control over heterochromatin formation and maintenance, which is essential for appropriate gene regulation and chromosomal integrity.

In recent years, multiple structural studies have demonstrated how histone ubiquitination regulates the activity of histone methyltransferases, significantly improving our understanding of the mechanisms underlying ubiquitination-methylation trans-crosstalk^57–59^. For instance, the monoubiquitination of histone H2B (H2BK120Ub)^60–64^ and H2B lysine 34 (H2BK34Ub)^46,65^ stimulate the methylation activity of Dot1L on histone H3 lysine 79 (H3K79)^66,67^ via enzyme positioning and nucleosome distortion-induced orientation restriction, respectively. H2BK120ub can promote histone H3 lysine 4 methylation (H3K4me) through allosteric COMPASS^68,69^/MLL^70–75^. Additionally, histone H2A lysine 119 monoubiquitination (H2AK119Ub) enhances the recruitment of PRC2 to increase the methylation level of histone H3 lysine 27 (H3K27)^76–78^. These studies revealed various mechanisms by which histone ubiquitination on the histone globular domain of nucleosomes establishes interchain crosstalk (i.e., in *trans* crosstalk) with methylation on nucleosomes. Stimulation of H3K9 methylation by H3K14Ub represents a form of histone intrachain *cis*-crosstalk that occurs on the histone tail and is independent of the histone globular domain. Our work provides structural insights into the ubiquitin-mediated crosstalk within the “histone tail network”, characterized by a synergistic mechanism that enhances both affinity and turnover of histone substrate.

It remains to be investigated whether the structural mechanism of H3K14Ub-stimulated Clr4 is conserved in metazoans. While Clr4 is the sole H3K9 methyltransferase found in yeast, humans possess various types of Suv39 family methyltransferases responsible for H3K9 methylation. Notably, evidence of mammalian Suv39 family methyltransferase regulation by ubiquitination has been reported, such as the essential role of K867 monoubiquitination in the SETDB1 SI-region for its enzymatic activity and function^79^. The ubiquitination site is close to the histone-binding groove, but this modification occurs on methyltransferases rather than the histone. Thus, further investigation to elucidate the mechanism of this crosstalk. Additionally, SUV39H2 (also known as KMT1B) was also found to be enzymatically activated by H3K14Ub^42^. The structure of SUV39H2 is very similar to that of Clr4, and especially, the hydrophobic and electrostatic interfaces with ubiquitin are both preserved: residue F189 of SUV39H2 may form the same critical hydrophobic interactions with ubiquitin as Clr4 F256, meanwhile in SUV38H2, the homology sites of Clr4 D280 and D281 are replaced by ^204^AE^205^, and Glu(E) likewise potentially form a salt bridge with Ub R42 **(Extended Data** Fig. 6e). In contrast, none of the corresponding Phe(F) or acidic amino acids are conserved in the SUV39H1 protein, which is consistent with the previously reported phenomenon that SUV39H1 does not sense ubiquitin. Interestingly, other acylation modifications have been reported for H3K14^80–82^. Among them, H3K14 acetylation (H3K14ac) is a marker of transcriptional activation^83^, functionally similar to H3K9 acetylation (H3K9ac), which opposes the crosstalk of H3K14Ub- H3K9me3 associated with the heterochromatin formation^84,85^. Exploring how these acylation modifications affect ubiquitination, heterochromatin formation, and maintenance of epigenetic traits may provide deeper insights into the intricate network of regulatory crosstalk mediated by Suv39 family methyltransferases.

Chemical protein synthesis provides a powerful tool for investigating the functions of complex histone modifications. By chemically synthesizing histone substrates with precise modifications, we can introduce modifications that are challenging to achieve using traditional biological methods (e.g., recombinant expression)^86^. In this work, this approach has allowed us to generate various samples for biochemical tests such as ubiquitinated peptides with native isopeptide bonds, structural analysis samples that mimic catalytic states (e.g., by replacing lysine with norleucine), and hypothesis verification samples such as those constructed with AEEA to artificially extend peptide bonds^46,87–90^. These synthetically derived samples enable controlled investigations of the effects of individual PTMs on enzymatic activity and chromatin structure, as well as the study of regulatory crosstalk among multiple modifications. The advancement of these synthetic strategies has significantly improved our understanding of histone modifications and their broader implications for epigenetic regulation.

## Methods

### Cloning and plasmid construction

The cDNA of *Schizosaccharomyces pombe* Clr4(Suv39h) was synthesized by GenScript Biotech (Nanjing, China) and subcloned into pGEX6p-1 vector downstream the GST-tag followed by an HRV3C cleavage site. Clr4 truncations including Clr4-KMT for crystallization and biochemistry studies, Clr4-ΔCD, and all the Clr4 mutants (F256A, D280AD281A, Y357A, D371A, F383A, D384A, D386A, H425A, F427A, F449AY451A, K455RK472R, K455AK472A, K455WK472W) were generated from the Clr4-pGEX6p-1 plasmid by standard site-directed PCR mutagenesis and homologous recombination, tagged by GST-HRV3C at N-terminus as well. The DNA sequences encoding the wild-type Ub and Ub_G76C_ were cloned into the pET22b vector without tag. All the Ub mutants used in this work (R42A, L8RH68A, L69AV70A, L71AL73A, R72AR74A) were generated from Ub_G76C_ pET22b plasmid. Genes of histone H2A, H2B, H3(all Cys to Ser), and H4, were cloned into pET22b respectively without tag. H3 mutants H3_K14C_, H3_K18C_, and H3_K23C_ were expressed in plasmid derived from H3-pET22b (all Cys to Ser). Meanwhile, the 147 bp Widom 601 DNA we user for nucleosome assembly along with histones was 64x repeat inserted into a vector. The sequence of 147 bp Widom 601 DNA: CTGGAGAATCCCGGTGCCGAGGCCGCTCAATTGGTCGTAGACAGCTCTAGCACCGCTTAAACGCACGTACGCGCTGTCCCCCGCGTTTTAACCGCCAAGGGGATTACTCCCTAGTCTCCAGGCACGTGTCAGATATATACATCCTGT At last, the genes of 3xFlag-tagged Clr4 or Clr4 mutants for *in vivo* assay were cloned into the pFA6a vector with *hph^+^* resistance from Danesh Moazed by homologous recombination, resulting in plasmids pFA6a-*hph*-*clr4*up-3xFlag-*clr4*variants-*clr4*down for *S. pombe* transfection.

### Protein expression and purification

Plasmids containing WT full-length Clr4 (or its variants F256A, D280AD281A, D280AD281R, Y357A, D371A, F383A, D386A, F449A/Y451A, F427A, K455AK472A, K455RK472R, K455WK472W) were transformed into BL21(DE3) Escherichia coli cells. The E. coli cells were grown in Luria broth (LB) medium containing 50 μg ml^−1^ ampicillin until an OD600 of 0.6. Cells were then induced by 0.4 mM isopropyl β-D-thiogalactopyranoside (IPTG) and 0.1 mM ZnCl_2_ at 16°C for 16 h. The cells were collected by centrifugation at 4,000 r.p.m. for 30 min and then lysed by sonication in the ice-cold lysis buffer (25 mM HEPES, pH 7.3, 200 mM NaCl, 10 μM ZnCl_2_, 1mM DTT and 1 mM phenylmethyl sulfonyl fluoride (PMSF)). After centrifugation at 14,000 r.p.m. for 30 min, the supernatant was incubated with glutathione-Sepharose 4B beads (GE Healthcare) for 2 h at 4°C. The proteins were washed with the elution buffer (25 mM HEPES, pH 7.3, 200 mM NaCl, 10 μM ZnCl_2_, 1 mM DTT, 30 mM GSH). The protein elution was incubated with PreScission Protease and dialyzed against 25 mM HEPES, pH 7.3, and 100 mM NaCl overnight at 4°C to remove the GST tag. The proteins were subsequently loaded into a Source 15S cation exchange column (GE Healthcare) equilibrated with the ion-exchange (IEX) Buffer A (25 mM HESPS, pH 7.3, 50 mM NaCl) and then eluted by a gradient of the IEX Buffer A and IEX Buffer B (50 mM HESPS, pH 7.3, 1 M NaCl). The eluted Clr4 proteins were further purified by a Superdex 75 10/300 GL size-exclusion column (GE Healthcare) pre-equilibrated in size-exclusion chromatography (SEC) buffer (25 mM HEPES, pH 7.5, 100 mM NaCl). The KMT domain of Clr4 (Clr4(192–490)) and its variants were purified as described above, except that a Source 15Q anion exchange column (GE Healthcare) was used to purify the proteins after the PreScission Protease digestion. The proteins were characterized by SDS-PAGE and visualized using a ChemiDoc MP Imaging System (Bio-Rad).

Ubiquitin and its variants (R42A, L8RH68A, L69V70A, L71L73A, R72R74A) were purified as previously described^46^ and the Gly76 of ubiquitin (or its variants) was mutated to Cys for the synthesis of disulfide bond linked ubiquitination substrates. In brief, plasmids for expression were transformed into BL21(DE3) competent cells. Cells were induced by 1 mM IPTG at 37°C overnight and were collected by centrifugation at 4,000 r.p.m. for 30 min. Cells were lysed by sonication in Ub lysis buffer (25 mM HEPES, pH 7.5, 150 mM NaCl). Cell lysates were supplemented with 1% perchloric acid and then clarified by centrifugation. The supernatant was dialyzed against 0.1% trifluoroacetic acid (TFA) overnight. The dialyzed mixture was further purified by a Source 15Q anion exchange column (GE Healthcare). The proteins were characterized by SDS-PAGE and visualized using a ChemiDoc MP Imaging System (Bio-Rad).

Histones including H2A, H2B, H3 (C96S/C110S), H4, and their mutants (H3K14C, H3K18C, H3K23C), were purified as previously described.^46^ In brief, histones were expressed inclusion body in BL21(DE3) Escherichia coli cells and were dissolved in the unfolding buffer (6 M guanidinium chloride (Gn·HCl), 20 mM Tris, pH 7.5, 1 mM DTT). After centrifugation at 14,000 r.p.m. for 30 min, the clarified supernatant was dialyzed against 0.1% TFA overnight. The dialyzed mixture was purified by RP-HPLC and characterized by LC-ESI-MS.

### Peptide synthesis

Histon H3 N-terminal peptides(H3(1-20), H3(1-20)_G12C_, H3(1-20)_K14C_, H3(1-20)_P16C_, H3(3-19)_K9NleK14C_, H3(1-20)_K9Nle_) and Ub(1-45)NHNH_2_ were synthesized using standard Fmoc solid-phase peptide synthesis (SPPS) protocols under standard microwave conditions (CEM Liberty Blue). Rink amino resin was used for the synthesis of carboxyl-terminal substrate, while hydrazine resin was used for the synthesis of hydrazine terminal substrate. The coupling cycle was programmed as previously reported^91^. In brief, 0.5 mg resin was used for coupling. Dimethylformamide (DMF) containing 10% piperidine and 0.1 M Oxyma was used for deprotection. 0.2 M Fmoc-protected amino acids, 0.14 M N,N’-diisopropylcarbodiimide (DIC), and 0.7 M Oxyma in DMF were used for amino acid coupling (10 min at 50°C for His and Cys, 90°C for other residues). After the coupling cycle, the resin was washed with dichloromethane (DCM) and treated with the cleavage mixture (1 mL thioanisole, 1 mL tri-isopropylsilane, 0.5 mL 1,2-ethanedithiol, 1 mL ddH_2_O, 16.5 mL trifluoroacetic acid) for 2h at 37°C. The crude peptides were obtained by precipitating the cleavage mixture with cold diethyl ether and then purified by RP-HPLC and characterized by LC-ESI-MS.

### Synthesis of isopeptide bond linked ubiquitination substrates

For the synthesis of isopeptide bond linked ubiquitination substrates H3(1-20)K14Ub (Ub-H3t), H3(1-20)_K9Nle_K14Ub, H3(1-20)K14AEEAUb (Ub-AEEA-H3t), H3(1-20)_K9Nle_K14AEEAUb, H3 terminal tail peptides (H3(1-20)K14Alloc, H3(1-20)_K9Nle_K14Alloc) were first synthesized by SPPS protocol, except that a building block of Fmoc-Lys(Alloc)-OH was used for coupling at position Lys14. After the coupling cycle, the peptide-resin was incubated with the BOC protection mixture (di-tertbutyl decarbonate: *N*,*N*-Diisopropylethylamine: DMF = 1:1:5) for 10 min to add the BOC protecting group at the N-terminal of the peptide. After protection, the resin was washed by DCM. And the Alloc protecting group was removed by incubating with Pd[P(C_6_H_5_)_3_]_4_ (60 mg) and Ph_3_SiH (600 μL) in 2 mL DCM at 37°C overnight. The peptide resin was washed with sodium diethyldithiocarbamate (200 mg) in 40 mL DMF to remove Pd. The peptide-resin was then used for SPPS and the ε-amino group on Lys14 can be further coupled with Ub(A46C-76). The products H3(1-20)K14Ub(A46C-76), and H3(1-20)_K9Nle_K14Ub(A46C-76) were purified from the resin by using the cleavage mixture as described above. H3(1-20)K14AEEAUb(A46C-76) was synthesized as described above, except that Fmoc-AEEA-OH was used as a building block in the subsequent coupling of the ε-amino group on Lys14 of H3(1-20).

Ala46 in Ub was mutated to Cys to enable hydrazide-based native chemical ligation^92^ for the next fragment Ub(1-45)NHNH_2_. In brief, Ub(1-45)NHNH_2_ (1 μM, 1.0 eq.) was dissolved in ligation buffer (6 M Gn·HCl, 100 mM NaH2PO4, pH 2.3) precooled to -15°C. Then, NaNO_2_ (10 μM, 10 eq.) was added, and the reaction was stirred for 30 min at -15°C to fully convert the hydrazide to acyl azide. The mixture was further treated with 4-mercaptophenylacetic acid (MPAA) (50 μM, 50 eq.) and the pH was adjusted to 5.0 to convert the acyl azide to thioester. Next, the N-terminal Cys peptides(1.2 μM, 1.2 eq.), were added into the mixture. Then, the pH was adjusted to 6.4 and stirred at 37°C overnight. Afer the reaction, 50 mM tris(2-carboxyethyl)phosphine (TCEP) was added to the mixture to reduce disulfide bonds and the pH was adjusted to 7.0. The products (H3(1- 20)K14Ub(A46C), H3(1-20)_K9Nle_K14Ub(A46C), H3(1-20)K14AEEAUb(A46C)) were then purified by RP-HPLC.

Then, the thiol group of Cys46 in Ub was removed by desulfurization reaction. The peptide was dissolved in a desulfurization buffer (6 M Gn·HCl, 100 mM Na_2_HPO_4_, pH 7.4). Subsequently, GSH (230 mg μM^−1^ peptide), 500 mM TCEP, and 20 mM VA-044 were dissolved in desulfurization buffer and added to the peptide solution. The pH was adjusted to 7.0 and the reaction solution was stirred at 37 °C overnight. The products were purified by RP-HPLC.

All the proteins (or peptides) involved in this section were characterized by LC-ESI-MS and lyophilized into powder for the next use.

To refold the Ub in H3 side chains, 6 mg freeze-dried products were dissolved in 6 M Gn·HCl, 100 mM HEPES, pH 7.4. And 250 μL 100 mM HEPES, pH 7.5 was added to reduce the concentration of Gn·HCl to 3M, and then the solution was dialyzed into a dialysis buffer (1 M Gn·HCl, 25 mM HEPES, pH 7.5). After 6 h, the dialysis buffer was replaced with no Gn·HCl buffer (25mM HEPES, pH 7.5, 150 Mm NaCl) for the next 6 h of dialysis. Finally, the product was purified by a Superdex 75 10/300 GL size-exclusion column (GE Healthcare) pre-equilibrated in SEC buffer (25 mM HEPES, pH 7.5, 100 mM NaCl).

### Synthesis of disulfide bond linked ubiquitination substrates

H3 peptides containing Cys mutation (H3(1-20)_G12C_, H3(1-20)_K14C_, H3(1-20)_P16C_, H3(3- 19)_K9NleK14C_, H3_K14C_, H3_K18C_, H3_K23C_) were concentrated to 10 mg/mL in disulfide ligation buffer (6 M Gn·HCl, 100 mM HEPES, pH 7.5). Then, 5,5’-dithiobis-(2-nitrobenzoic acid) (DTNB, Sigma Aldrich; 3.0 eq., from 0.1 M stock solution prepared in25 mM HEPES, pH 7.5, 100 mM NaCl) was added. The reaction was placed at room temperature for 1 h and quenched by adding HCl to adjust the pH to 1.0. All the H3-S-TNB products were purified by RP-HPLC and lyophilized into powder for the next use.

The product H3-S-TNB (10mg/mL, 1.0 eq.) and UbG76C (10mg/mL, 1.0 eq., or its variants) were separately dissolved in the disulfide ligation buffer. Next, the UbG76C solution was titrated to the H3-S-TNB solution and the pH was adjusted to 7.0. The reaction was bathed at 30°C water for 15 min and quenched by adding HCl to adjust the pH to 1.0. All the H3-Ub_G76C_ products were purified by RP-HPLC and lyophilized into powder for the next use.

The refolding of the Ub modified on H3t was performed as described above. The full-length H3-Ub_G76C_ (H3_K14C_-Ub_G76C_, H3_K18C_-Ub_G76C_, H3_K23C_-Ub_G76C_) will be refolded in the process of octamer reconstitution and assembled into NCP_H3K14Ub_, NCP_H3K18Ub_, and NCP_H3K23Ub_.

### Crystallization and structure determination

For the crystal structure of KMT-Ub-H3, the Clr4 KMT, H3(3-19)_K9NleK14CUb_, and S-adenosylmethionine (SAM) were incubated in the ratio of 1:1.05:5 at a final KMT concentration of 24.5 mg/mL for 6 hours, 4 °C. Crystallization was under 291K with sitting-drop vapor diffusion. The complex was first crystallized in the reservoir buffer of 20% PEG1000, 100 mM Imidazole, 200 mM CaAc_2_, pH 7.0 (1:1). The initial crystals were broken down by ultrasonication in the reservoir buffer and then utilized as seeds in the crystallization system consisting of 0.5 μL protein complex solution and 0.5 μL 18-20% PEG1000, 100 mM Imidazole, 200 mM CaAc2, pH 7.0. The KMT-H3 complex sample was obtained by incubating the KMT, H3(3-19)_K9NleK14C_, and SAM at 1:10:10 with 20 mg/mL KMT. Satisfying crystals were also grown under 291K with sitting-drop vapor diffusion, in the reservoir solution containing 0.1 M Magnesium chloride hexahydrate, 0.1 M MES, pH 6.0, 8 % w/v PEG 6000. All harvested crystals were cryo-protected in the solution with 10 % glycerol in the respective reservoir solution, and preserved in liquid nitrogen.

Both the diffraction of the crystals of KMT-Ub-H3 and KMT-H3 complex was collected under the wavelength of 0.979176 Å and temperature of 100 K at the SSRF BEAMLINE BL02U1 of Shanghai Synchrotron Radiation Facility. Raw diffraction data KMT-Ub-H3 and KMT-H3 complex were processed and scaled by Aimless 0.7.4^93^ and HKL2000^94^, respectively. We used the previously reported apo WT Clr4 KMT structure (PDB 6BP4) and Ub crystal structure (PDB 1UBQ) for molecular replacement phasing in PHENIX 1.19.2_4158^95^. The model building was visualized and operated by COOT^96^ and collection and refinement statistics of our structures are presented in Extended Data Table 1.

### Enzyme/substrate mutation methyltransferase assay

The methyltransferase activity of WT Clr4 or its variants (F256A, D280AD281A, Y357A, D371A, F383A, D384A, D386A, H425A, F427A, F449AY451A, K455AK472A, K455RK472R, K455WK472W) was detected by the MTase-Glo™ Methyltransferase Assay (Promega Corporation). In brief, 40 μL 2X enzyme solution (100 nM/400 nM WT Clr4 or its variants (400 nM Clr4 was used when the substrate was unmodified H3t, while 100 nM Clr4 for Ub-H3t), 80 μM SAM, 10 μM ZnCl_2_, diluted by SEC buffer and 40 μL 2X substrate solution (80 μM H3t or 4 μM Ub-H3t, diluted by SEC buffer were incubated at 30°C and then were mixed as the start of the reaction. The mixed solution was taken 8 μL at 0 min, 10 min, 20 min, or 1 h, and subsequently added into 384-Well solid white assay plate pre-filled with 2 μL 0.5% TFA at each time point to stop the reaction. Each well of the assay plate was added 2 μL 6X MTase-Glo™ Reagent and incubated at room temperature for 30 min. Then each well of the assay plate was added 12 μL 2X MTase-Glo™ Detection Solution and incubated at room temperature for 30 min. After the incubation, the luminescence was measured by using a plate-reading luminometer (BioTek). The influence of substrate variants in the Clr4 methyltransferase activity was detected in a similar protocol to that for Clr4 variants, except that only the WT Clr4 was used in 2X enzyme solution and the substrate variants (Ub-H3t mutants, H3tG_C_12- Ub, H3tP_C_16-Ub, Ub-AEEA-H3t, NCP (2 μM), NCP_H3K14Ub_ (2 μM), NCP_H3K18Ub_ (2 μM), NCP_H3K23Ub_ (2 μM)) were used in 2X substrate solution.

The production of SAH was calculated by a standard curve using serial dilutions of SAH. The protocol was described as follows. 2 μM SAH was serially diluted to half the concentration each time in the range of 2 µM to 0 µM (2 µM, 1 µM, 0.5 µM, 0.25 µM, 0.125 µM, 0.063 µM, 0.031 µM, 0 µM, 8 μL each sample). Consistent with the above processing, 2 μL 0.5% TFA, 2 μL 6X MTase- Glo™ Reagent, and 12 μL 2X MTase-Glo™ Detection Solution were added and incubated for the corresponding time. The luminescence was measured by the plate-reading luminometer. The standard curve was generated by plotting luminescence (Y-axis) against SAH concentration (X-axis) so that the linear equation was established.

Automethylated Clr4 was performed as in the previous report with minor modifications^36^. In brief, 2.4 μM Clr4 or its variants were dialyzed in the HMT buffer (50 mM Tris-HCl, pH 8, 20 mM KCl, 10 mM MgCl_2_, 1 mM DTT, 0.02% Triton, 5% glycerol, 1 mM PMSF) supplemented with 250 μM SAM for 4 h at room temperature with mild agitation and then were dialyzed in the SEC buffer for 6 h. After incubation, the methyltransferase activity of Clr4 was analyzed as described above.

### Determination of Michaelis constant

Michaelis constant of WT Clr4 (or its variants Clr4-H425A, Clr4-F427A) with its substrates (Ub-H3t or H3t) was also determined by using MTase-Glo™ Methyltransferase Assay. Ub-H3t was serially diluted in SEC buffer with the concentration in a range of 16 μM to 0 µM (16 µM, 8 µM, 4 µM, 2 µM, 1 µM, 0.5 µM, 0.25 µM, 0.125 µM, 0 µM, 55 μL each sample) to prepare a series of substrate solution. 2X enzyme solution (20 nM Clr4 or its variants, 160 μM SAM, 2 μM ZnCl_2_, diluted by SEC buffer) and 2X substrate solution were incubated at 30°C. Then each substrate solution was added with 55 μL 2X enzyme solution to start the reaction. The reaction solution was taken for 8 μL at 6 min and then subsequently added into a 384-Well solid white assay plate pre-filled with 2 μL 0.5% TFA to stop the reaction. Consistent with the methyltransferase assay, 2 μL 6X MTase-Glo™ Reagent, and 12 μL 2X MTase-Glo™ Detection Solution were then added and incubated for the corresponding time. When the substrate was H3t, the experiment process was the same as that of Ub-H3t, except that the enzyme concentration of 2X enzyme solution was changed to 200 nM and the concentration of 2X substrate was changed to a range of 128 μM to 0 µM and the sampling time was changed to 60 min.

The luminescence was measured by the plate-reading luminometer. The production of SAH was also calculated by an SAH standard curve as described in the methyltransferase assay. Data fitting and analyses were performed using GraphPad Prism 9.0.

### Western blot for methyltransferase activity

NCP (WT NCP, NCP_H3K14Ub_; 1 µM, 5 eq.), SAM (100 µM, 500 eq.), ZnCl2 (10 µM; 50 eq.), GSSG (500 µM) were diluted by SEC buffer and incubated at 30℃. Reactions were commenced by the addition of Clr4 (200 nM, 1 eq.). 1 µL was removed from each reaction at 10 min, 20 min, 60 min, 120 min, 200 min, 300 min and then boiled with LDS sample buffer (Thermo Fisher Scientific) at 95℃ for 5 min. Boiled samples were resolved on a 4−12% gradient SDS-PAGE gel at 160 volts for 25 min. Gels were transferred onto PVDF membranes by eBlot^TM^ L1 (GenScript Biotech). Membranes were blocked for 40 min with QuickBlock blocking buffer (Beyotime) and incubated with corresponding primary antibodies at 4℃ overnight. After incubation, membranes were washed with TBST buffer (50 mM Tris-HCl, pH 7.6, 150 mM NaCl, and 0.1% (v/v) Tween 20) for 5 min for 5 times. And then membranes were incubated with HRP-conjugated secondary antibody for 45 min at room temperature followed by a 5-minute TBST buffer wash 5 times. Finally, membranes were treated with an ECL Substrate (Clarity Max^TM^ Western, Bio-Rad) and visualized using a ChemiDoc MP Imaging System (Bio-Rad). Antibodies were diluted by antibody dilution buffer (Primary & Secondary Antibody Diluent for WB, YEASEN). Antibody information is listed as follows: rabbit polyclonal anti-H3K9me1 antibody (39888, Active Motif, 1/2000 dilution), mouse monoclonal anti- H3K9me2 antibody (ab1220, Abcam, 1/4000 dilution), rabbit polyclonal anti-H3K9me3 antibody (ab8898, Abcam, 1/5000 dilution), rabbit polyclonal anti-H2A antibody (ab18255, Abcam, 1/1000 dilution), HRP* Goat Anti Mouse IgG(H+L) (RS0001, Immunoway, 1/10000 dilution), HRP* Goat Anti Rabbit IgG(H+L) (RS0002, Immunoway, 1/10000 dilution).

### Isothermal titration calorimetry analysis (ITC)

The isothermal titration calorimetry data were collected using a MicroCal PEAQ-ITC (Malven Pananlytical). Clr4^KMT^ and ligands (Ub, Ub-H3t, H3(1-20)_K14C_-Ub_G76C_, H3(1-20)_K14C_-Ub_G76C_(R42A), H3(1-20)_K14C_-Ub_G76C_(L7173A), H3(1-20)_K14C_-Ub_G76C_(L8RH68A)) were buffer-exchanged into dialyzed overnight against ITC buffer (25 mM Na_2_HPO_4_ pH 7.5, 100 mM NaCl) prior to experiments. For the experiments, 300 μL 20 μM substrates solution in the sample cell was titrated with 200 μM Clr4^KMT^ solution through 19 injections (2.0 μl each) at 25 °C and 750 r.p.m. stirring speed. Data fitting and analyses were performed using PEAQ-ITC instrument client software.

### Thermo-shift assay

The thermos-shift of Clr4-KMT banding with substrates was evaluated using an UNcle all-in-one biologics stability screening platform (Unchained Labs). Cl4-KMT (60 μM, 1 eq.), SAM (600 μM, 10 eq.), and a corresponding substrate (60 μM, 1 eq.) were mixed in SEC buffer and loaded on the UNcle. The temperature started at 20 [and ended at 95 [with an increasing rate of 0.3-1 [/min. The fluorescence spectra of each sample was analyzed by UNcle Analysis software. The melting temperature was deter-mined by the first derivative of the barycentric mean (BCM) of fluorescence intensity with respect to temperature (dBCM/dT).

### Reconstitution of octamers and nucleosomes

Nucleosomes containing unmodified H3, H3_K14C_-Ub_G76C_, H3_K18C_-Ub_G76C_, H3_K23C_-Ub_G76C_ were reconstituted as previous report^97^. In brief, the four core histones H2A, H2B, H3(C96S/C110S), and H4 were mixed at a stoichiometry of 1.1:1.1:1:1 and dialyzed against the refolding buffer (10 mM HEPES, pH 7.5, 2 M NaCl, 1 mM EDTA) for 18 h with buffer changed every 6 h. The desired octamers were purified by using a Superdex 200 10/300 GL size-exclusion column (GE Healthcare) pre-equilibrated in refolding buffer. Next, the nucleosomes were reconstituted by mixing the octamers and 147-bp Widom 601 DNA at a stoichiometry of 1:1.1 in the refolding buffer. And the NaCl concentration was gradient reduced to below 200 mM by adding HE buffer (10 mM HEPES, pH 7.5, 1 mM EDTA) with the use of a peristaltic pump at 4 [overnight. To further reduce the NaCl concentration, the mixture was dialyzed against the HE buffer at 4 [overnight. Assembled nucleosomes were purified by using anion exchange on a DEAE-5PW column (TSKgel) pre-equilibrated in HE buffer. Desired nucleosomes were eluted by a gradient of HE Buffer and 1 M NaCl HE buffer. Octamers and nucleosomes were characterized by SDS-PAGE (GeneScript).

### Electrophoretic mobility shift assay (EMSA)

WT full-length Clr4 was prepared as a twofold dilution series in SEC buffer (25 mM HEPES, pH 7.5, 100 mM NaCl) in a range of 8 μM to 0.625 μM. Next, 5 μL Clr4 with indicated concentrations was mixed with 5 μL 200 nM NCP (or NCP_H3K14Ub_, NCP_H3K18Ub_) diluted by the SEC buffer. The mixed samples were incubated for 10 min at 4°C. 5 μL Native gel loading (0.15% (v/v) bromophenol blue, 50% (v/v)) glycerol, 0.25% (v/v) BSA) was then added to each sample and mixed well. Subsequently, 5 μL of each sample was resolved by 4.5% native–PAGE gels running in a pre-cooled 0.5 × TBE buffer (40 mM Tris-HCl, pH 8.3, 45 mM boric acid, and 1 mM EDTA) at 150 V for 45 min in ice water. Gels were stained with 0.01% (v/v) SYBR-Gold dye (Thermo Fisher Scientific) and visualized using a ChemiDoc MP Imaging System (Bio-Rad).

### Strains construction and Silencing assays

All yeast mutants used in this article (except Clr4 knockout strain) were transformed from the gift strain (SPY7283) obtained from Moazed lab, and the transformation method was the same as reported before^36^. Among them, the Clr4 knockout strain (SPY7305) was also obtained from the Moazed lab. The mutant constructions were confirmed by PCR amplification and sequencing to determine their correctness.

Silencing experiments were performed similarly to the previously reported article^36^. Cells were harvested at a final concentration of 1 × 10^7^ cells/mL, washed once with sterile water, resuspended in 150 μL sterile water, and then serially diluted 10-fold. 3 μL of each dilution was dropped onto the appropriate growth medium (YE). The plates were incubated in a 32 °C incubator for 2 days and then left at 4 °C for 36 hours to enhance red pigmentation before imaging to assess the silencing of the *ade6^+^* reporter gene.

## Acknowledgments

We thank the National Key R&D Program of China (no.2022YFC3401500, for L. Liu), National Natural Science Foundation of China (nos.22137005, 92253302, 22227810, T2488301 for L. Liu). L. Liu was also supported by the XPLORER prize and the New Cornerstone Investigator Program. We thank the Tsinghua University Branch of China National Protein Science Facility, Tsinghua University, Beijing, for the device support of crystal screening (mosquito, SPT Labtech) and checking (XtaLAB FR-X, Rigaku) and the guidance about the crystallization experiment from Min Li (National Protein Science Facility, Tsinghua University). The data collection was supported by the Shanghai Synchrotron Radiation Facility (BL02U1). Protein MS analysis was performed by Meng Han in Proteomics Facility at Technology Center for Protein Sciences, Tsinghua University. We thank the laboratory of Danesh Moazed (Department of Cell Biology, Harvard Medical School, Howard Hughes Medical Institute, Boston 02115, U.S.) for supporting our yeast growth experiments.

## Author Contributions

L. Liu, H. Ai, and Q. Qu. supervised the project. M. Sun, Y. Du, H. Ai, and L. Liu proposed the idea and analyzed the results. M. Sun, Y. Du, X. Wu, H. Ai, Q. Qu, and L. Liu designed the experiments. M. Sun, Y. Du, H. Ai, and Z. Li cloned the plasmids, expressed the proteins (Clr4 and histones, including mutants), and reconstituted the nucleosomes. Y. Du prepared the crystal samples and collected the crystal data. Y. Du solved the structures of the Clr4 KMT domain bound to the ubiquitinated or unmodified H3 peptide. M. Sun, Y. Du, and Z. Li performed the methyltransferase assay. Y. Du conducted the electrophoretic mobility shift assay (EMSA). Y. Du completed the thermal shift assay. Y. Du and Z. Li conducted isothermal titration calorimetry analysis (ITC). Y. Du, Z. Li, M. Sun, and X. Wu synthesized the peptides or ubiquitinated peptides for biochemical exploration. M. Sun and Y. Du performed strain construction and yeast growth assay. M. Sun, Y. Du, 3. H. Ai, and Z. Li wrote and revised the manuscript. M. Sun, Y. Du, H. Ai, Z. Li, Q. Qu, X. Wu, and L. Liu, read and analyzed the manuscript.

## Competing interests

The authors declare no competing interests.

## Data availability

Crystal structures of the KMT-Ub-H3 and KMT-H3 complex have been published in the Protein Data Bank (PDB ID code 9ISZ and 9IT4 respectively). The apo WT Clr4 KMT structure and Ub crystal structure are available under PDB accession code 6BOX, 6BP4 and 1UBQ.

---

**Extended Data Fig. 1.**
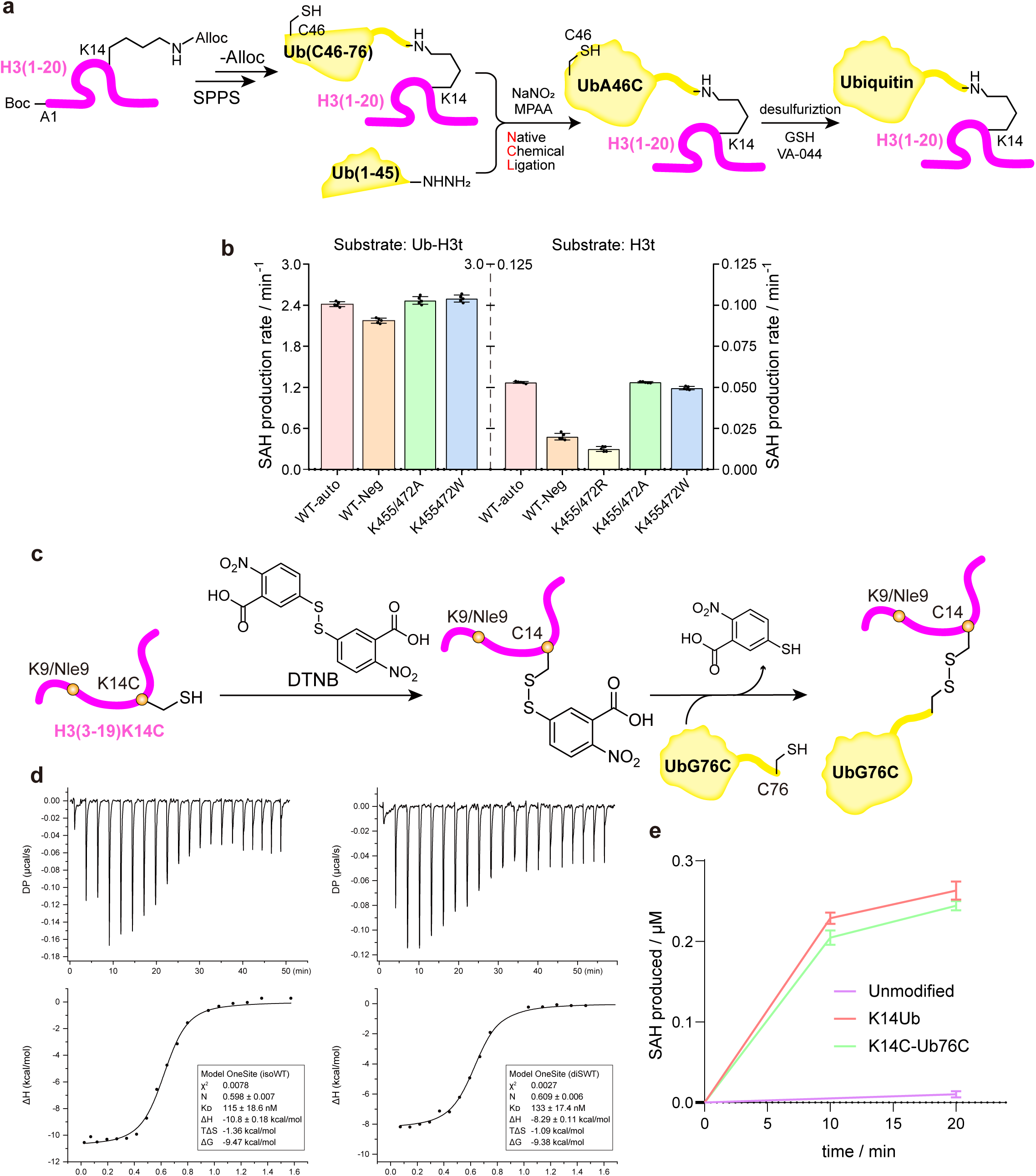
Chemical synthesis of substrates and related biochemical tests. **a.** Scheme of chemical synthesis of the K14-ubiquitinated H3(1-20) (Ub-H3t), with the ubiquitin moiety indicated in yellow and the histone moiety in magentas. This strategy is used for all isopeptide bond-linked ubiquitinated substrates hereinafter. **b.** The average methyltransferring rate per micromolar enzyme during the methylation of Ub-H3t or H3t mediated by WT Clr4, automethylated Clr4, and Clr4 replaced K455/K472 by R/A/W. The average rate was calculated directly from the endpoint quantity of SAH detected by the chemiluminescence activity assay (n=5, error bars indicate SD). **c.** Scheme of chemical synthesis of the H3(3-19)_K9Nle-K14C_-Ub_G76C_. This strategy is used for all disulfide bond-linked ubiquitinated substrates hereinafter. **d.** ITC experiment performed with WT Clr4 KMT and Ub-H3t (left) or H3(1-20)_K14C_-Ub_G76C_ (right). The top is the raw titration calorimetry curve and the lower is the fitted curve and the fitted thermodynamic constant in each picture. **e.** Histone methyltransferase activity assay of WT Clr4-KMT catalyzing methylation of H3t, Ub-H3t, and H3(1-20)_K14C_-Ub_G76C_ (n=3, error bars indicate SD).

**Extended Data Fig. 2.**
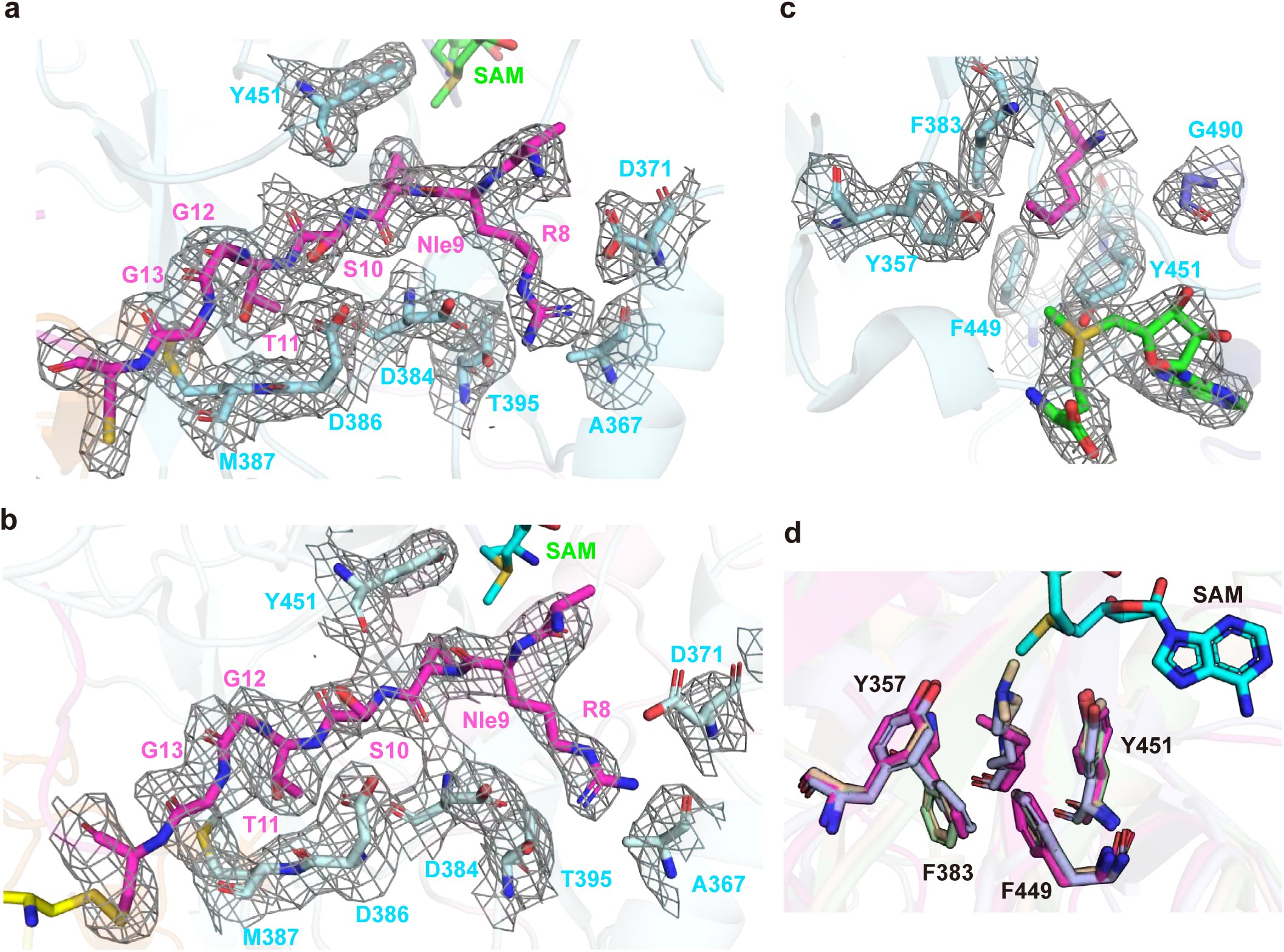
**Atomic details of interactions between Clr4 KMT and H3 N-terminal peptide**. **a, b.** Residues involved in the interactions between H3 N-terminal tail and Clr4 in **a)** KMT- H3 (Clr4 KMT bind to H3(3-19)_K9Nle-K14C_) and **b)** KMT-Ub-H3 (Clr4 KMT bind to H3(3-19)_K9Nle-K14C_-Ub_G76C_). H3 is in magentas and residues of Clr4 are represented by cyan sticks, while in KMT- H3 are green. The 2Fo–Fc electron density of residues of interest is shown in gray mesh contoured at 1.0 σ, and 0.8 σ, respectively. **c.** Details of SAM and K_Nle_9 binding with the Clr4 catalytic site in KMT-Ub-H3 structure. SAM is in green and the 2Fo–Fc electron density of residues of interest is shown in gray mesh contoured at 1.0 σ. **d.** Comparison of conformation of the catalytic site among structures of Suv39 proteins binding with H3 N-terminal and SAM. Clr4 and SAM in our structure are in magentas and cyan, while DIM5 (PDB: 1PEG) is in purple, G9a (PDB: 5JIY) is in pale green, and GLP1 (PDB: 2RFI) is in wheat.

**Extended Data Fig. 3.**
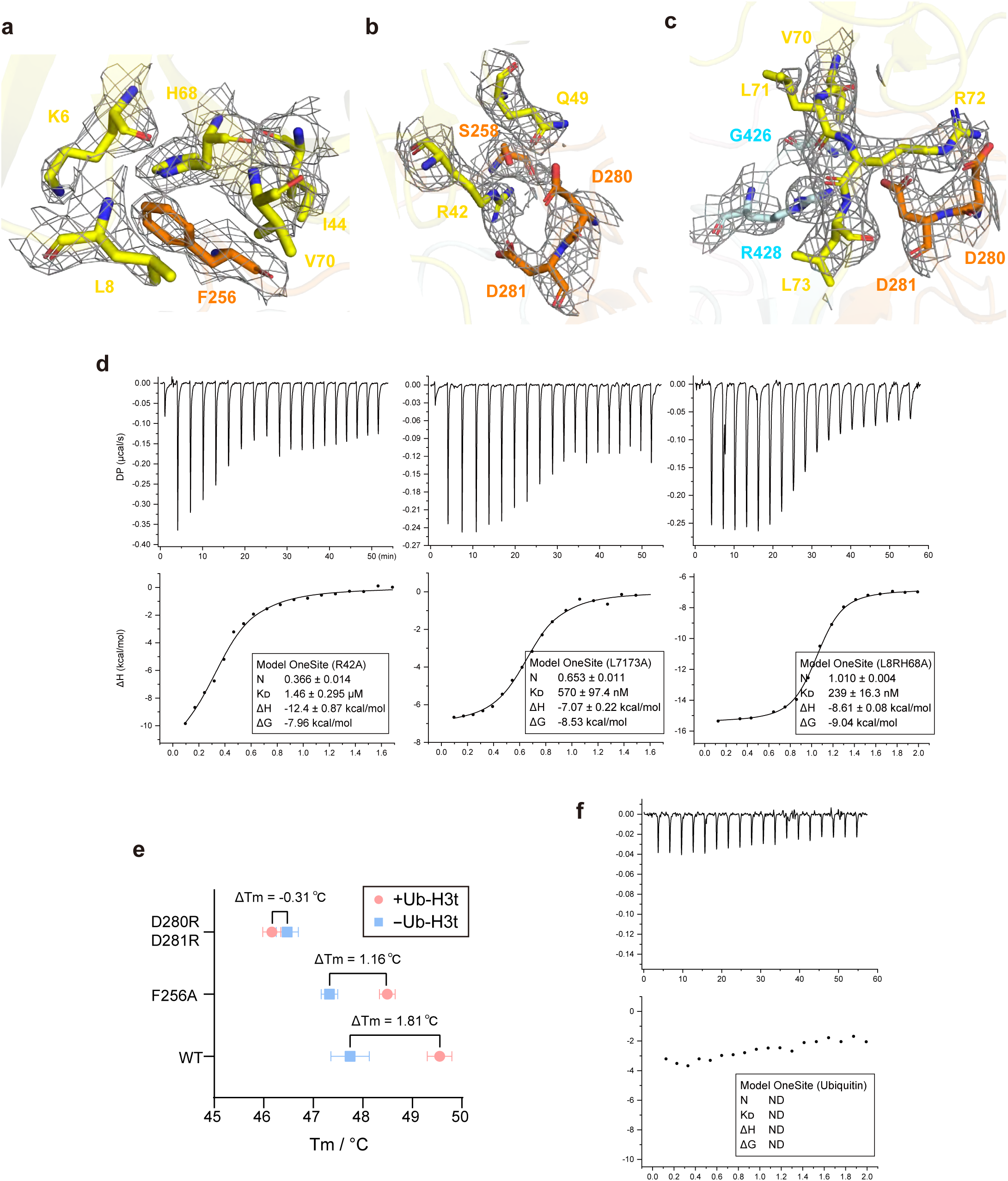
Interfaces between ubiquitin and Clr4 KMT. a, b, **c.** Key residues at the **a)** interface I, **b)** interface II, and **c)** interface III between Ub and Clr4 in KMT-Ub-H3 structure. Residues of Clr4 are represented by orange and cyan sticks, while Ub is colored by yellow. **d.** ITC experiment performed with WT Clr4 KMT and Ub-H3t variants with Ub mutation R42A (left), L71A/L73A (middle), and L8RH68A (right). The top is the raw titration calorimetry curve and the lower is the fitted curve and the fitted thermodynamic constant. **e.** Melting temperature *Tm* measured by the thermoshift assay based on WT Clr4 or Clr4 D280A/D281A, F256A mutants. Δ*Tm* before (blue) and after (pink) binding to substrate Ub-H3t are shown on the top of each group (n=3, error bars indicate SD). **g.** ITC experiment performed with WT Clr4 KMT and free ubiquitin.

**Extended Data Fig. 4.**
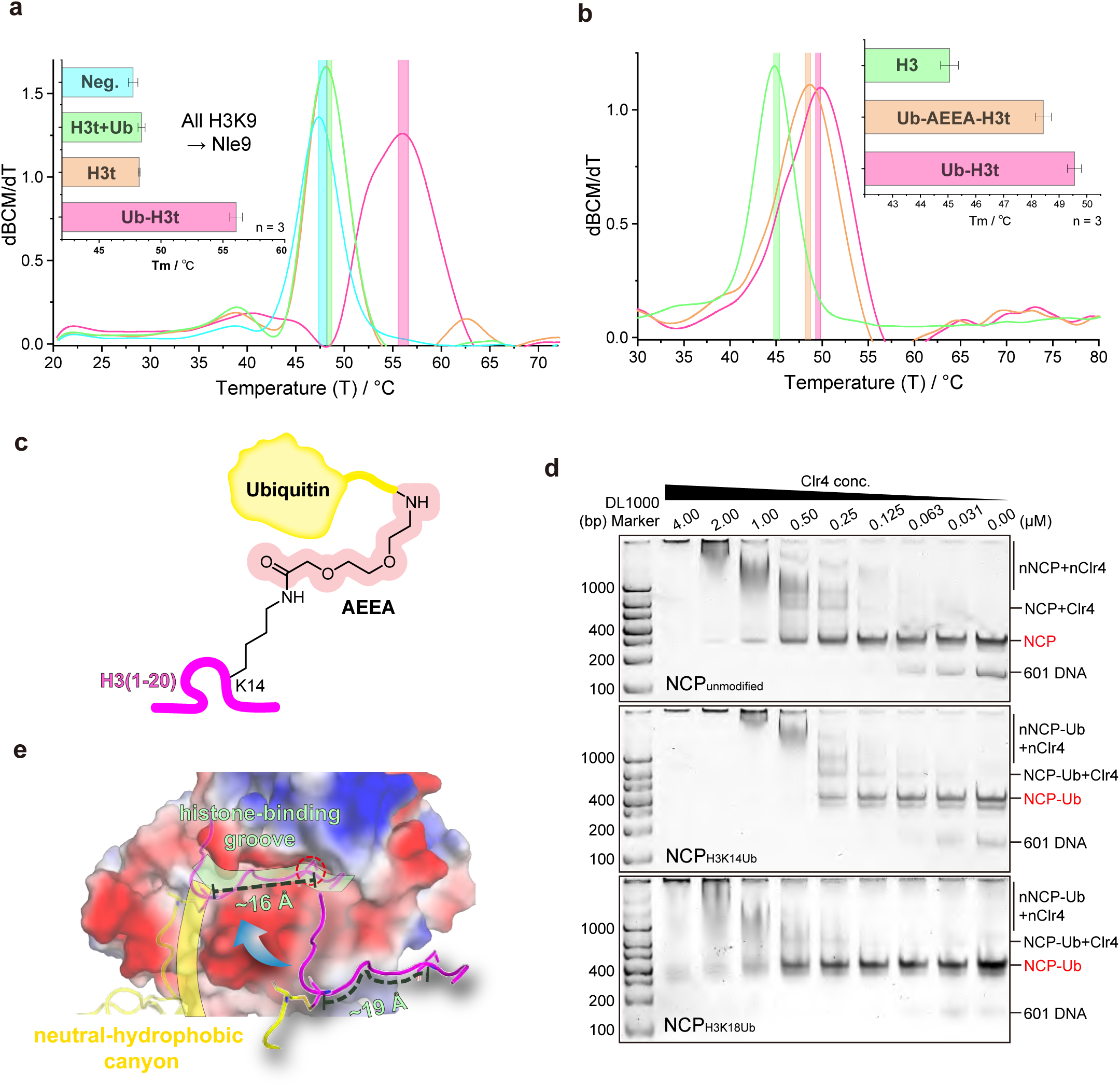
Biochemical experiment about H3-Ub junction. **a.** Thermo-shift assay working on Clr4 KMT binding with Ub-H3t(K_Nle_9), H3t(K_Nle_9), H3t(K_Nle_9) + free Ub. The larger picture represents the melting curve, the abscissa is the temperature (*T*) while the ordinate is the first derivative of the barycentric mean (BCM) of fluorescence intensity with respect to temperature (dBCM/d*T*). The top left smaller picture is the bar diagram of the melting temperature (*Tm*) of each sample (n=3, related to Fig. 4b). **b.** Thermo-shift assay performed with Clr4 KMT and H3t, Ub-H3t, Ub-AEEA-H3t, respectively (n=3, related to Fig. 4c). **c.** The molecular structure of the Ub-AEEA- H3t, the AEEA part is highlighted by the red contour. **d.** EMSA used to examine the binding affinity of Clr4 to different types of NCPs: the top is unmodified NCP, the middle is H3K14-ubiquitinated NCP, and the lower is H3K18-ubiquitinated NCP. The concentration of Clr4 in the mixture stepped up from right to left. **e.** Comparison of peptide chain lengths with surface groove lengths. Clr4 is represented by the electrostatic surface, H3 is in magentas and the ubiquitin is in yellow. Histone-binding groove is marked by a green patch and the neutral-hydrophobic canyon is marked by a yellow patch, while the catalytic site is highlighted by a red dash circle.

**Extended Data Fig. 5.**
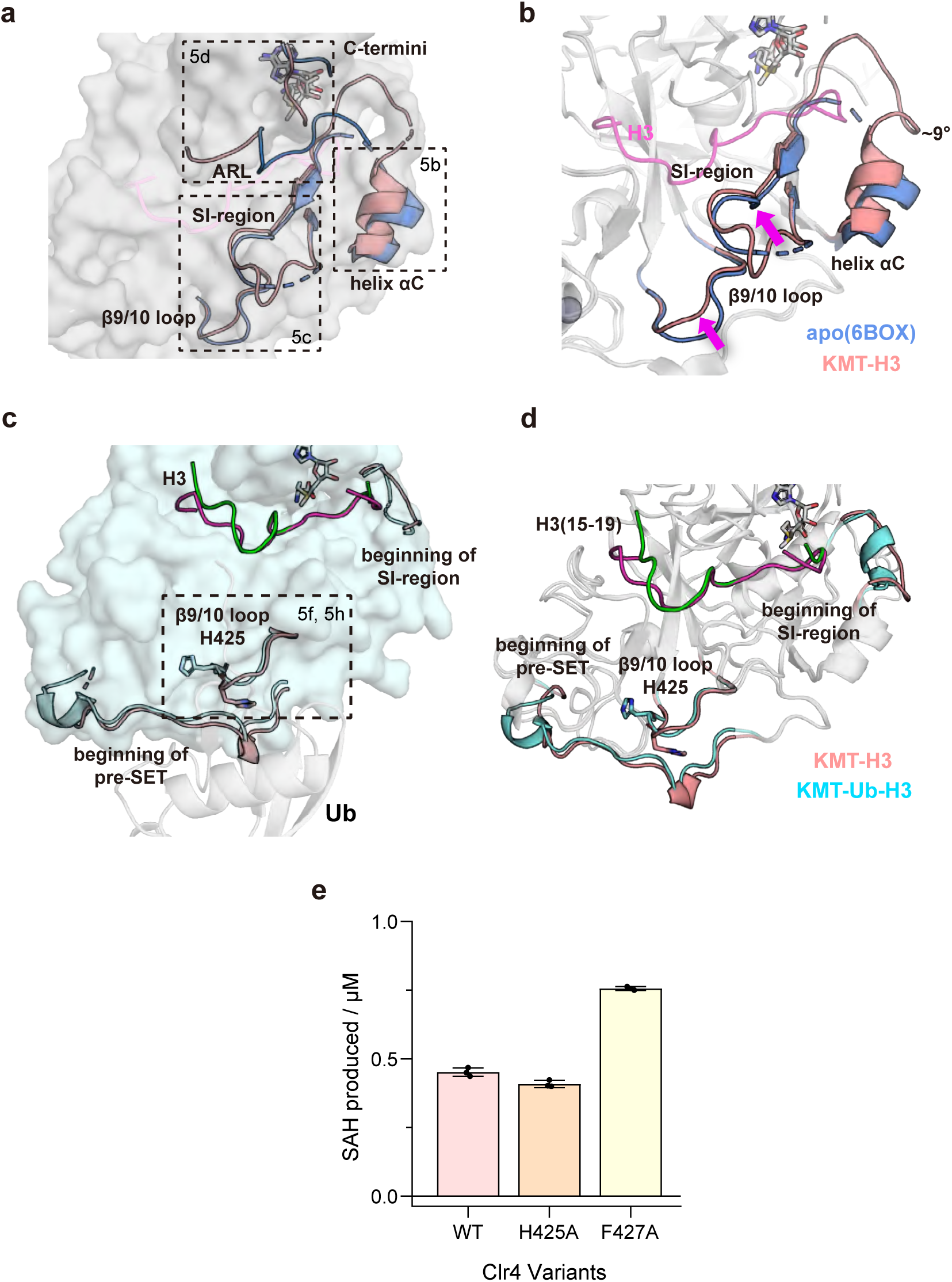
Conformational changes during H3t and Ub-H3t binding. **a.** Overview of conformational changes between the Clr4 KMT apo structure (6BOX) and our KMT-H3 structure. Two KMT structures are superposed and shown as a gray surface, structural elements undergoing conformational changes in the KMT-H3 structure are colored in pink and those in the apo form structure are colored in blue. **b.** Conformation changes of helix αC, SI region, and β9/10 loop between apo structure and our KMT-H3 structure. **c, d.** Conformational differences between Clr4 in KMT-H3 structure and KMT-Ub-H3 structure. Two KMT structures are superposed and shown as **c)** cyan surface or **d)** gray cartoon. For KMT-H3, Clr4 is colored in pink and H3 is in magentas, and for KMT-Ub-H3, Clr4 is colored in cyan and H3 is in green. **e.** Chemiluminescence activity assay of WT Clr4, Clr4 H425A, and Clr4 F427A working on unmodified substrate H3t (n=3, error bars indicate SD).

**Extended Data Fig. 6.**
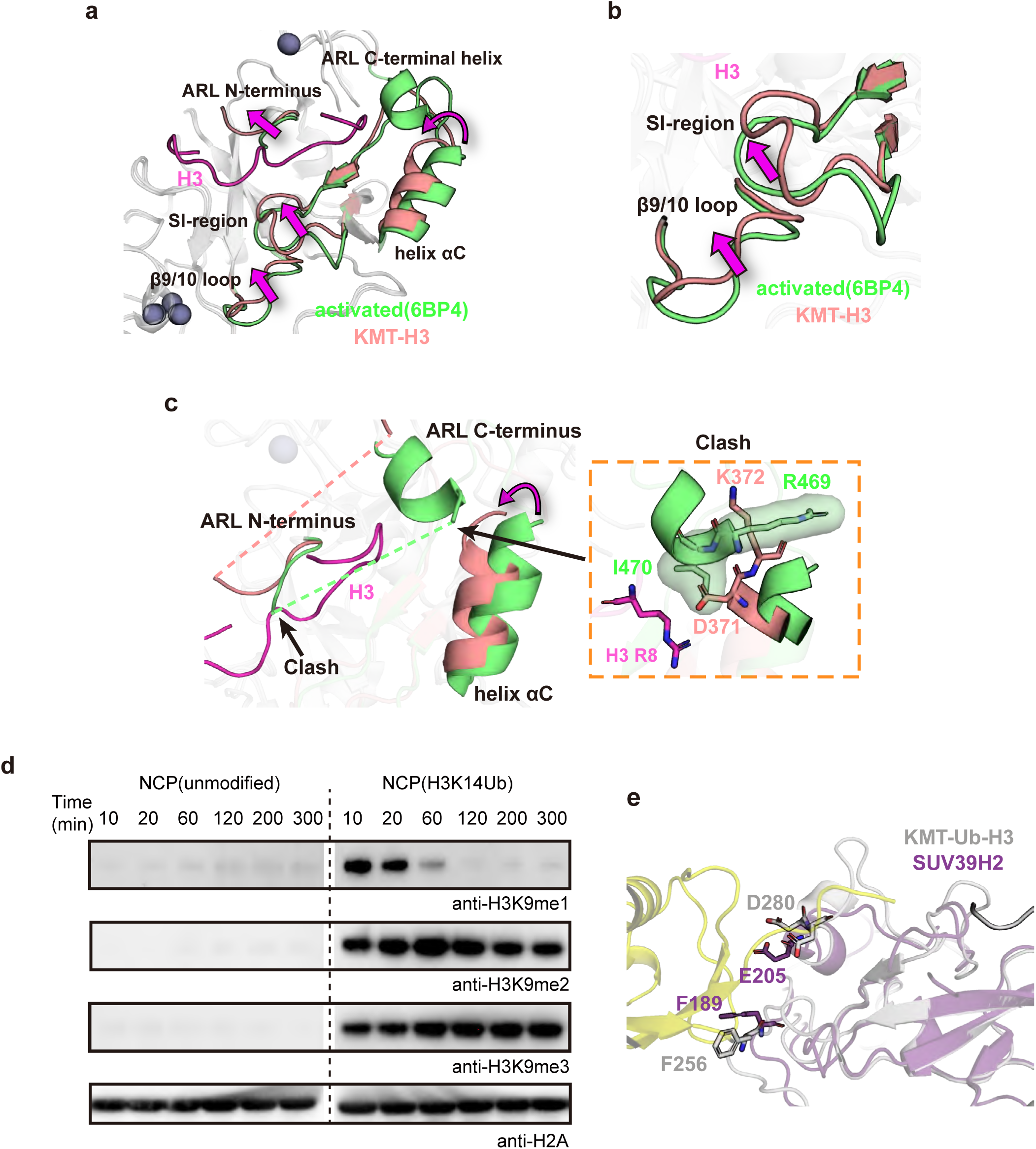
Different allosteric effects mediated by Clr4 automethylation and H3K14Ub. **a.** Collection of conformation changes from automethylated “activate state” (PDB: 6BP4) to the KMT-H3 structure. KMT domain in 6BP4 and KMT-H3 are superposed, and structural elements undergoing allostery are labeled and colored in green (6BP4) or pink (KMT-H3). **b.** Close- up views of allostery of SI region and β9/10 loop. **c.** The spatial conflict between the helix αC of KMT-H3 and the ARL C-terminal helix of 6BP4, as well as the clash between H3 and ARL N- terminal residues of 6BP4. The detail of the clash between ARL C-terminal helix I470 and helix αC D371 is shown on the right. **d.** Clr4 in *vitro* methylation with unmodified (left) and ubiquitinated (right) nucleosomes as substrates, respectively. H3K9me1, H3K9me2, and H3K9me3 produced in the reaction were detected by respective antibodies. H2A was used as an internal control for the system. **e.** Alignment of the structure of SUV39H2 (PDB: 6P0R, in purple) and KMT-Ub-H3 (in grey and yellow).

**Extended Data Table 1.**
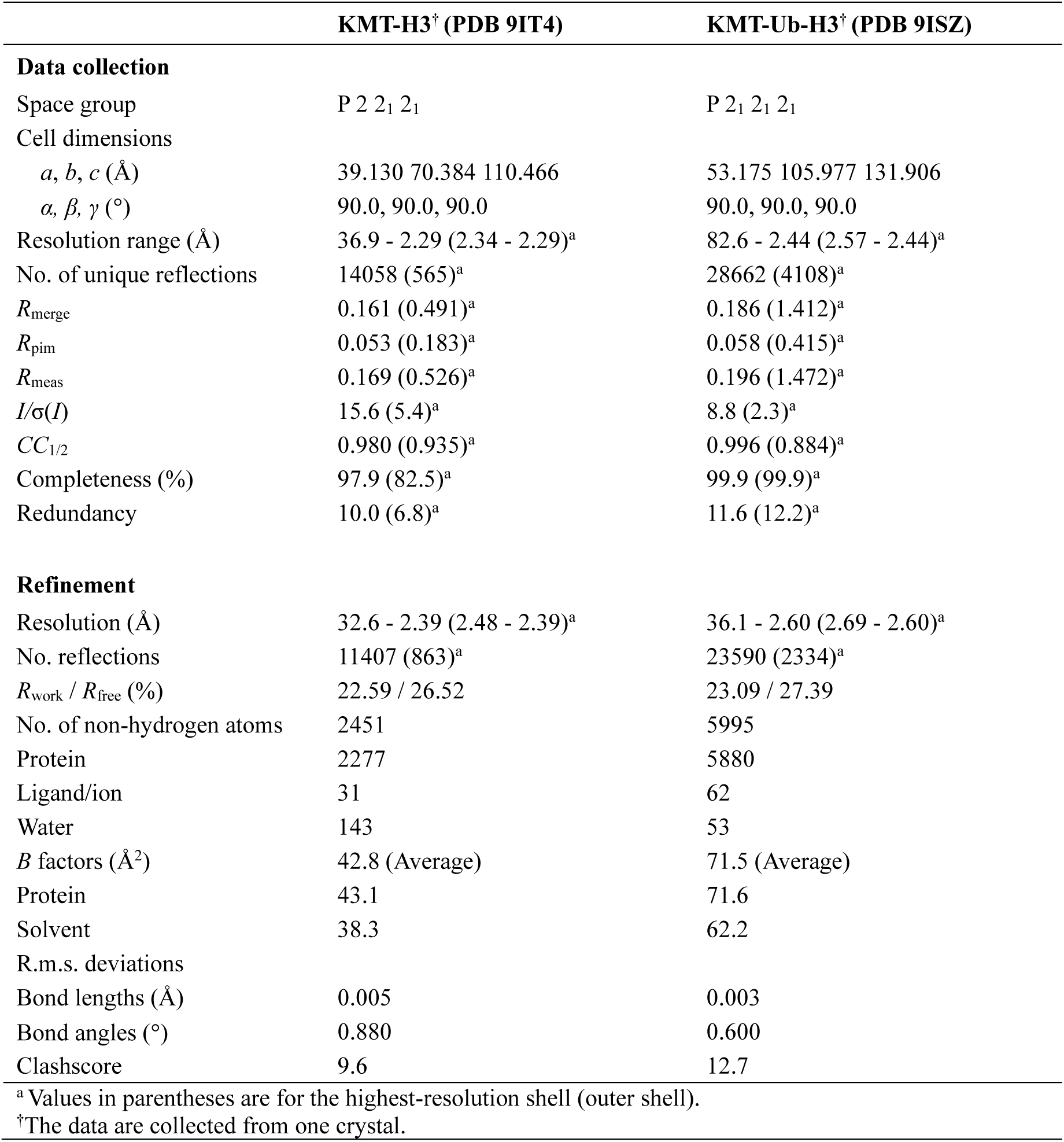
| Crystallographic data collection and refinement statistics.

## Notes

### Competing Interest Statement

The authors have declared no competing interest.

