## Supplementary Figures for "Mechanistic Insights into the Stimulation of the Histone H3K9 Methyltransferase Clr4 by Proximal H3K14 Ubiquitination"

### Supplementary Figure 1

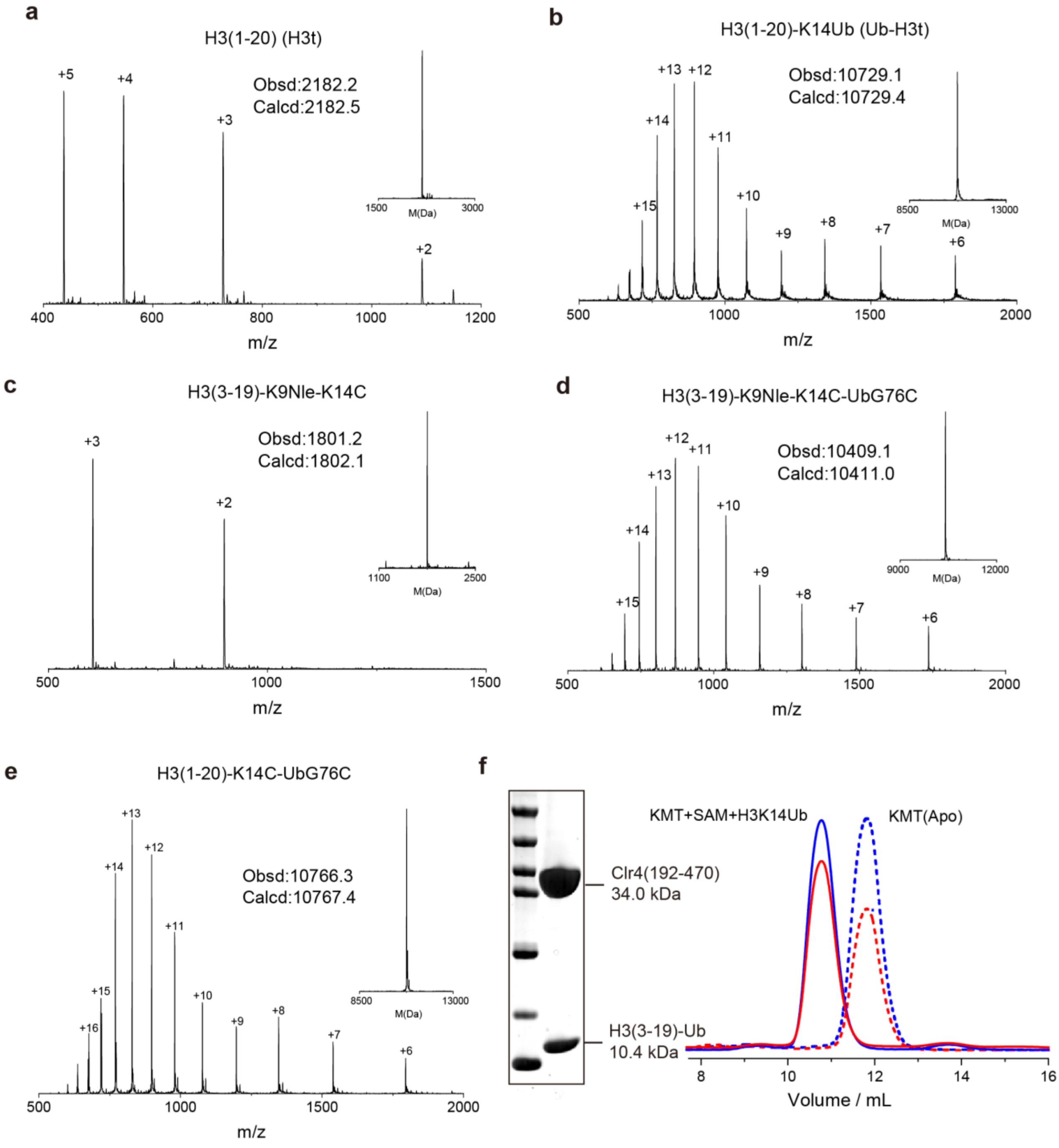

**Supplementary Fig. 1 | Property characterization of ingredient and protein samples in reconstitutions and crystallology. a-e.** Representative ESI-MS traces of **a)** H3(1-20) (H3t), **b)** H3(1-20)-K14Ub (Ub-H3t), **c)** H3(3-19)$_{K9Nle\text{-}K14C}$, **d)** H3(3-19)$_{K9Nle\text{-}K14C}$-Ub$_{G76C}$, and **e)** H3(1-20)$_{K14C}$-Ub$_{G76C}$. The observed (obs.) and calculated (calc.) molecular weights are marked, and the corresponding deconvoluted spectrum is shown on the top right. **f.** Size-exclusion chromatogram of Clr4 KMT and the KMT-Ub-H3 complex, the peak shift after complex assembly is shown, the blue line represents the 280 nm absorbance and the red line represents the 260 nm absorbance. The SDS-PAGE of the KMT-Ub-H3 complex for crystallization is shown on the left.

### Supplementary Figure 2

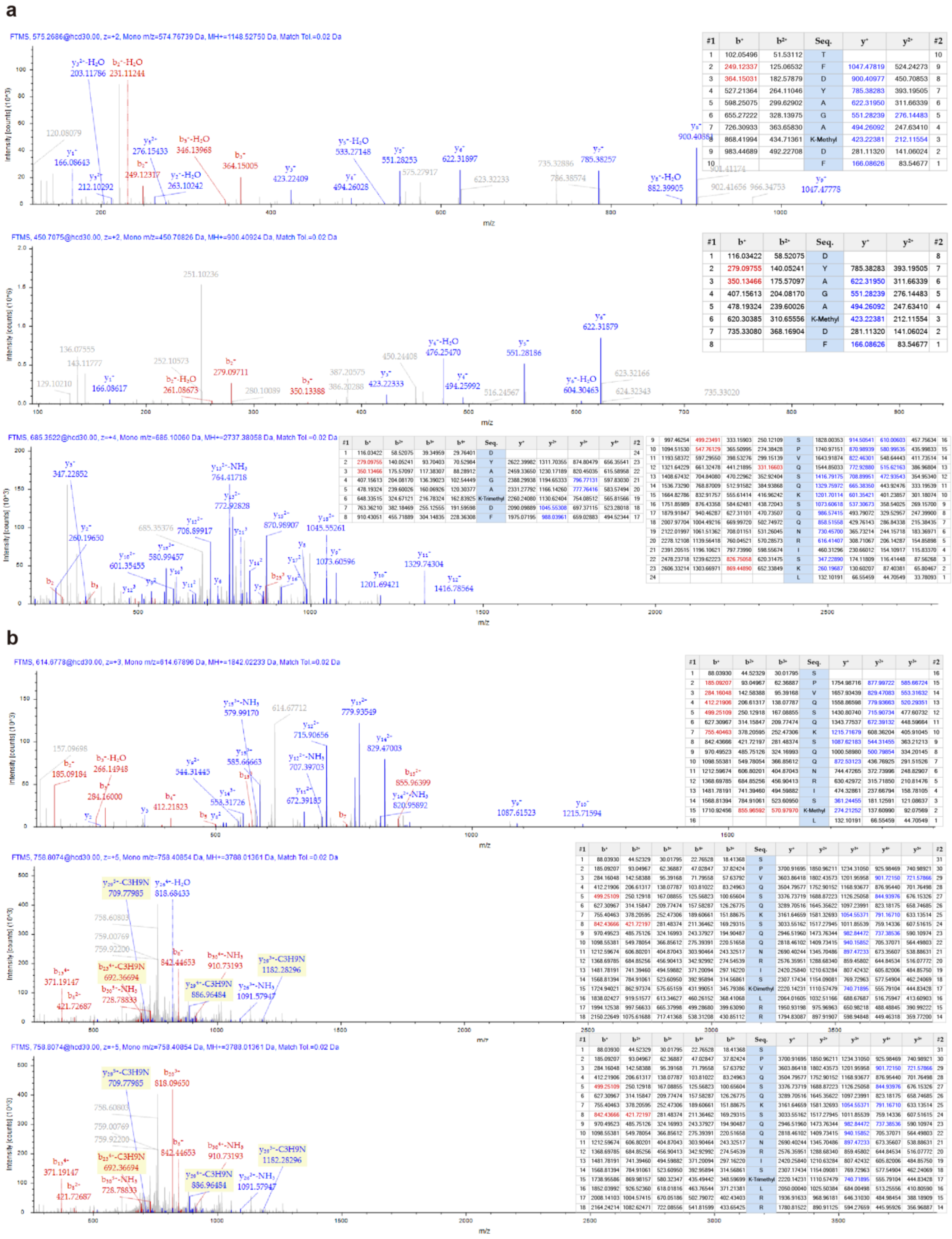

**Supplementary Fig. 2 | Identification of post-translational modifications on Clr4 by LC-MS/MS**. **a, b.** MS/MS spectrum confirming the automethylation on Clr4 **a)** K455, and **b)** K472. The peptide fragment sequence and corresponding theoretical y/b cation masses were listed in the table on the right. The experiment was repeated two times with chymotrypsin digestion and resulted in similar results.

### Supplementary Figure 3

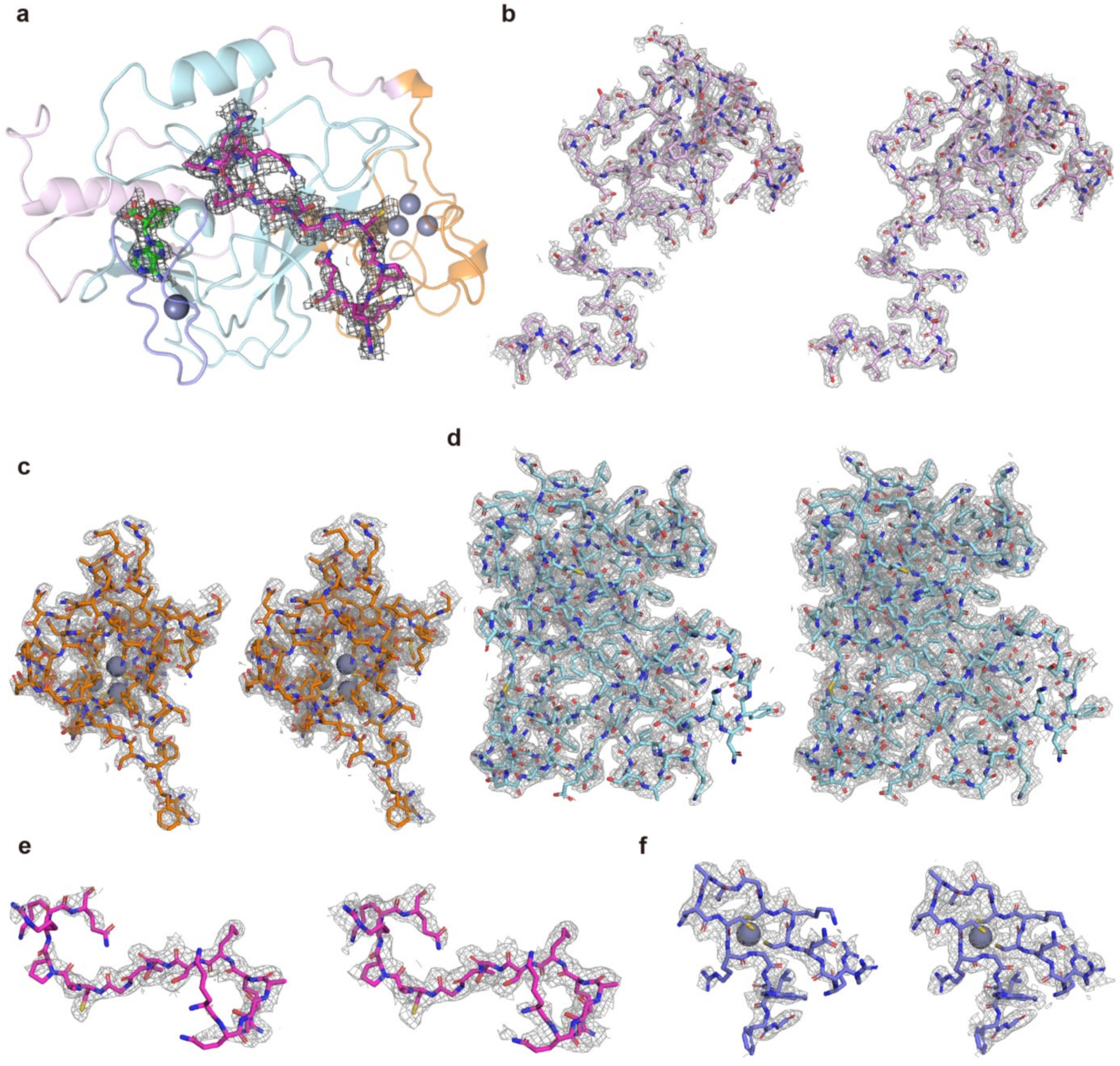


**Supplementary Fig. 3 | Structure and corresponding election density map of KMT-H3 complex.** **a.** Global structure of KMT-H3 complex. The 2Fo–Fc electron density of the peptide H3(3-19)$_{\text{K9Nle-K14C}}$ and SAM is shown in grey mesh contoured at 1.0 σ. The Clr4 subdomain NT is colored in pink, pre-SET is colored in orange, SET is colored in cyan, and post-SET is colored in purple. **b-f.** Structure and corresponding election density map of **b)** NT, **c)** pre-SET, **d)** SET, **f)** post-SET subdomain and **e)** H3(3-19)$_{\text{K9Nle-K14C}}$. The mesh is contoured at 1.0 σ (left) and 0.6 σ (right).

### Supplementary Figure 4

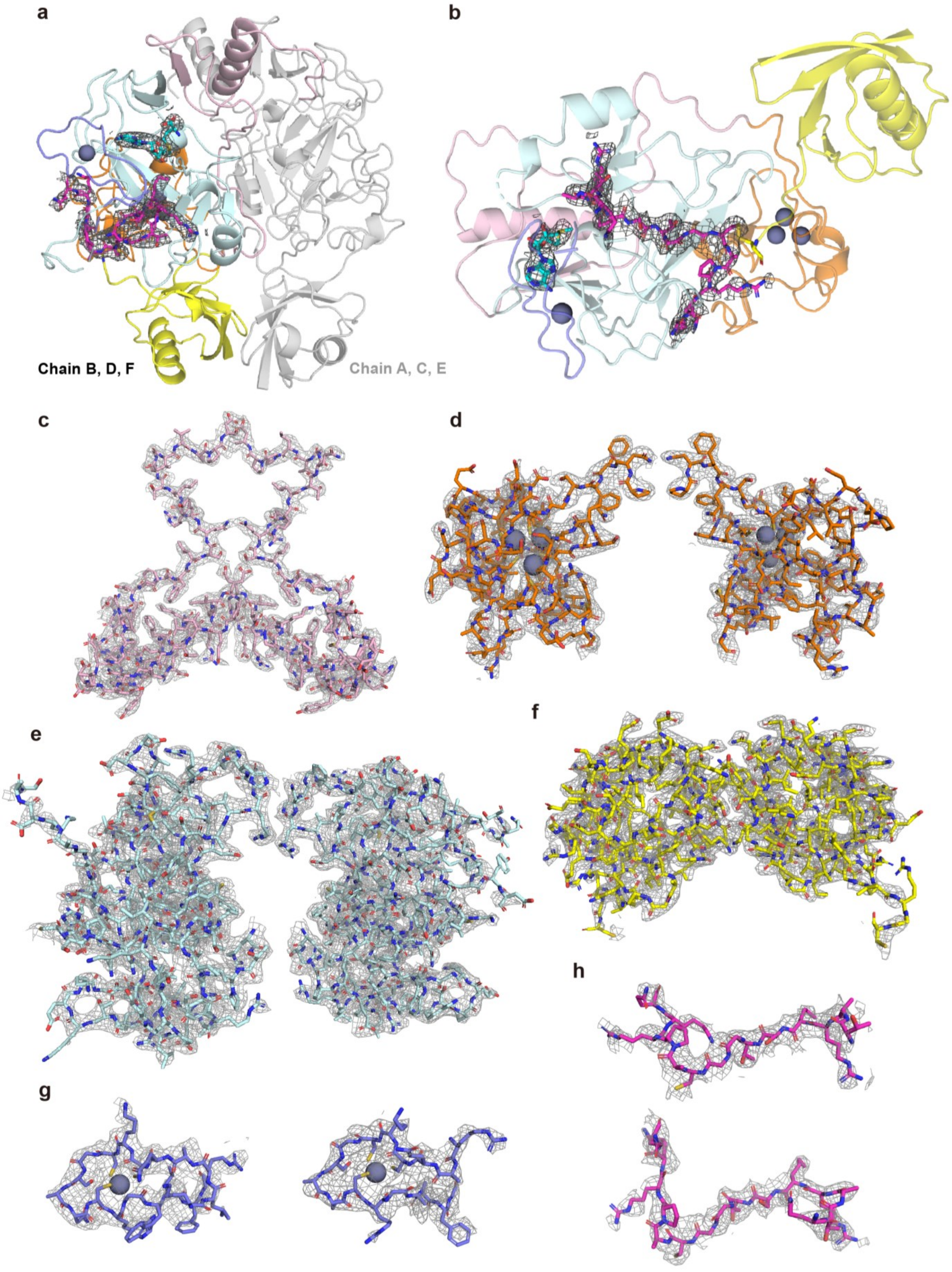

**Supplementary Fig. 4 | Structure and corresponding election density map of KMT-Ub-H3 complex.** **a.** Global structure of dimerized KMT-Ub-H3 complex. The 2Fo–Fc electron density of the peptide H3(3-19)$_{K9Nle-K14C}$-Ub$_{G76C}$ and SAM is shown in grey mesh contoured at 1.0 σ. Chain A, C, and E were in grey and chain B (Clr4) and F (H3) are colored the same as supplementary Fig. 3a, chain D (Ub) is colored in yellow. The Clr4 subdomain NT is colored in pink, pre-SET is colored in orange, SET is colored in cyan, and post-SET is colored in purple. **b.** Structure of NCS unit of chain B, D, F. **c-h.** Structure and corresponding election density map of **c)** NT, **d)** pre-SET, **e)** SET, **g)** post-SET subdomain, and **f)** ubiquitin, **h)** H3(3-19)$_{K9Nle-K14C}$. The mesh is contoured at 0.6 σ.

### Supplementary Figure 5

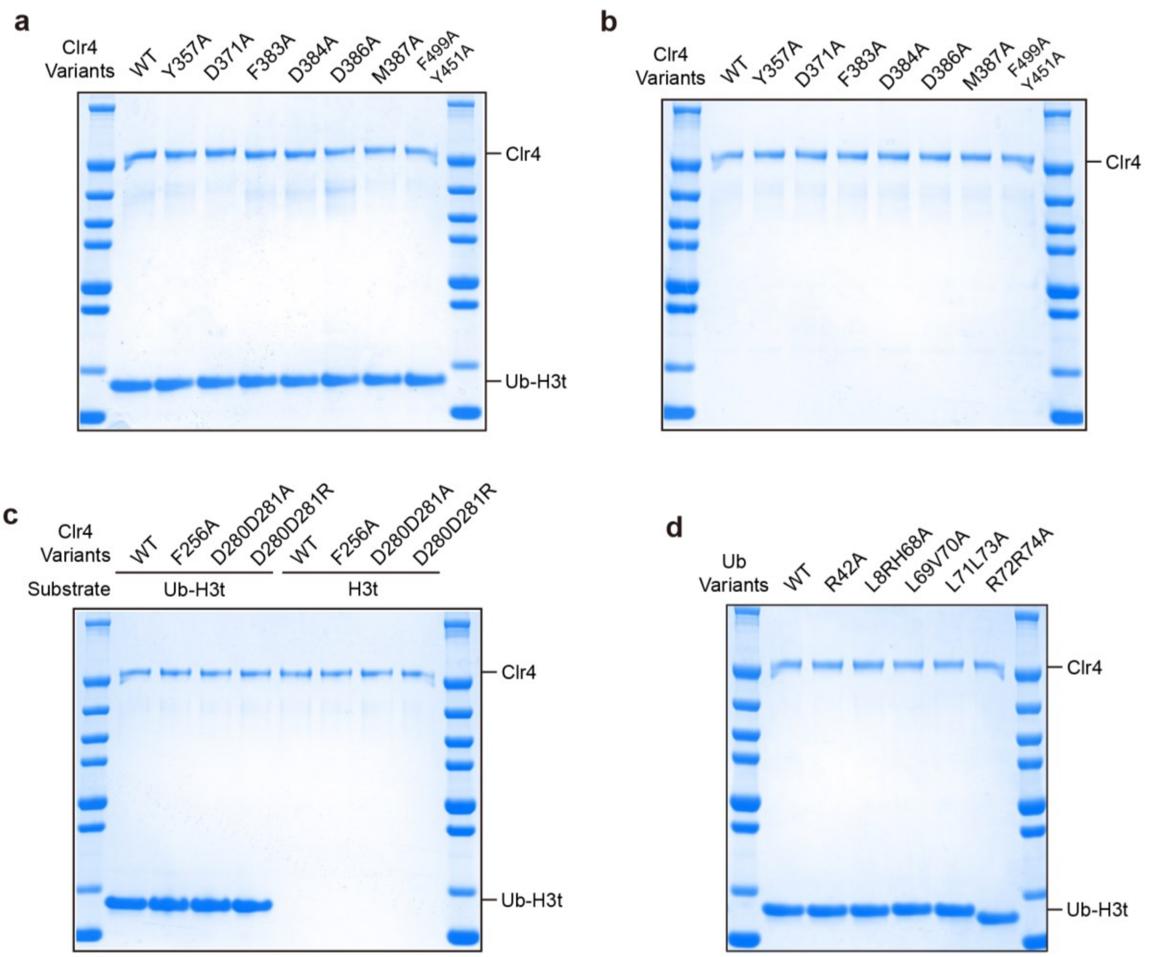


**Supplementary Fig. 5 | Methyltransferring reaction system with mutated protein samples. a.** SDS-PAGE of the reaction system of different Clr4 variants accommodating mutations in the histone-binding groove catalyzing Ub-H3t methylation. The SDS-PAGE sample is prepared from the reaction mixture without buffer dilution, as the actual final concentration of Clr4 is too low to be visible. **b.** SDS-PAGE of the reaction system of different Clr4 variants accommodating mutations in the histone-binding groove catalyzing H3t methylation. **c.** SDS-PAGE of the reaction system of different Clr4 variants accommodating mutations in Clr4-Ub interfaces catalyzing Ub-H3t and H3t methylation. **d.** SDS-PAGE of the reaction system of WT Clr4 variants catalyzing Ub-H3t variants methylation.

### Supplementary Figure 6

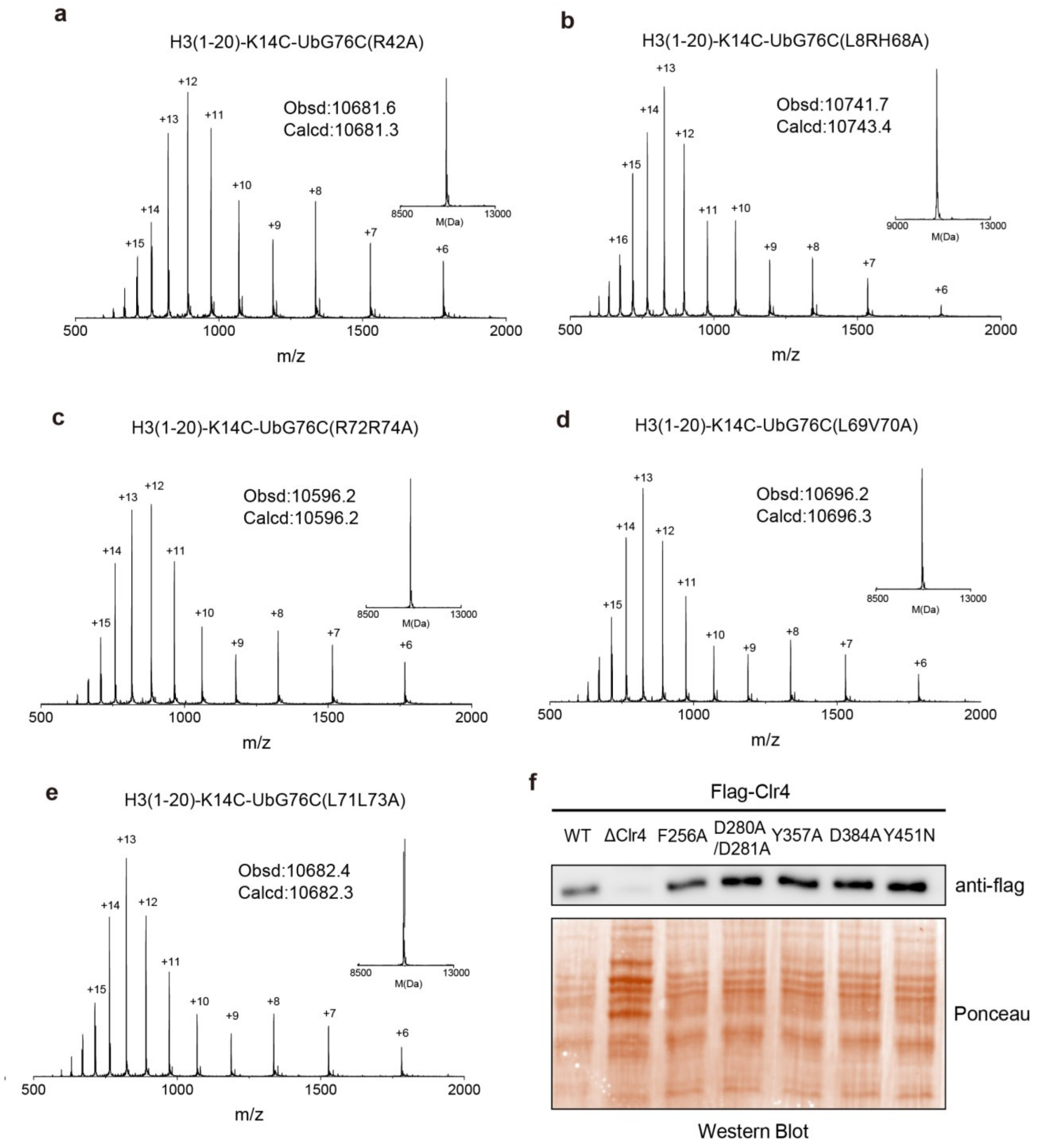


**Supplementary Fig. 6 | Validation of substrate samples and fission yeast strain. a-e.** Representative ESI-MS traces of **a)** Ub(R42A)-H3t, **b)** Ub(L8R/H68A)-H3t, **c)** Ub(R42A/R74A)-H3t, **d)** Ub(L69A/V70A)-H3t, and **e)** Ub(L71A/L73A)-H3t. **f.** Western blot of expression of Flag-Clr4 or mutated Flag-Clr4 in different yeast mutants. Clr4 is detected by FLAG antibody, and the internal reference was performed by ponceau staining.

### Supplementary Figure 7

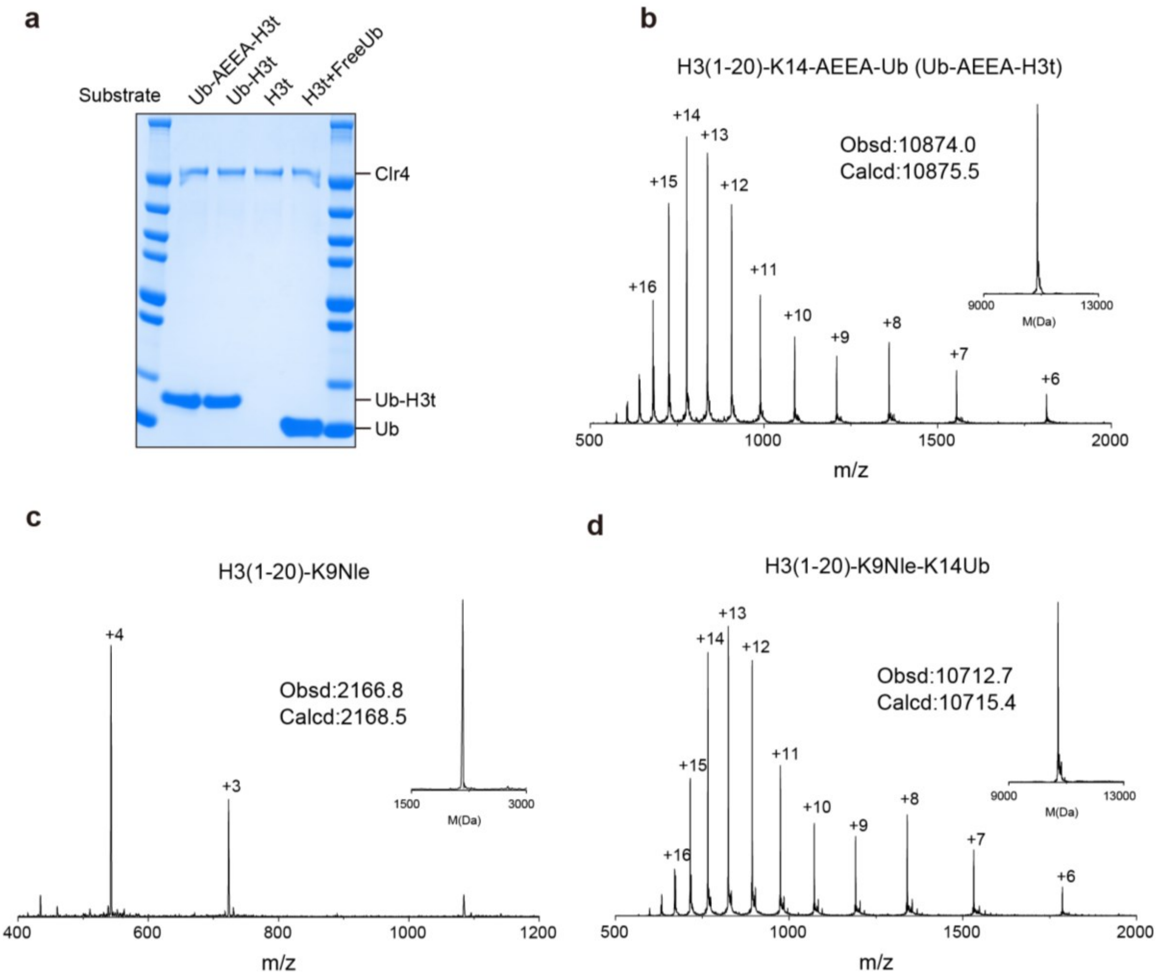


**Supplementary Fig. 7 | Property characterization of substrate samples for exploration of essentiality of H3K14-Ub linkage. a.** SDS-PAGE of the reaction system of WT Clr4 variants catalyzing Ub-H3t, Ub-AEEA-H3t, and H3t methylation. **b-d.** Representative ESI-MS traces of **b)** Ub-AEEA-H3t, **c)** H3(1-20)$_{K9Nle}$, **d)** H3(1-20)$_{K9Nle}$-K14Ub.

### Supplementary Figure 8

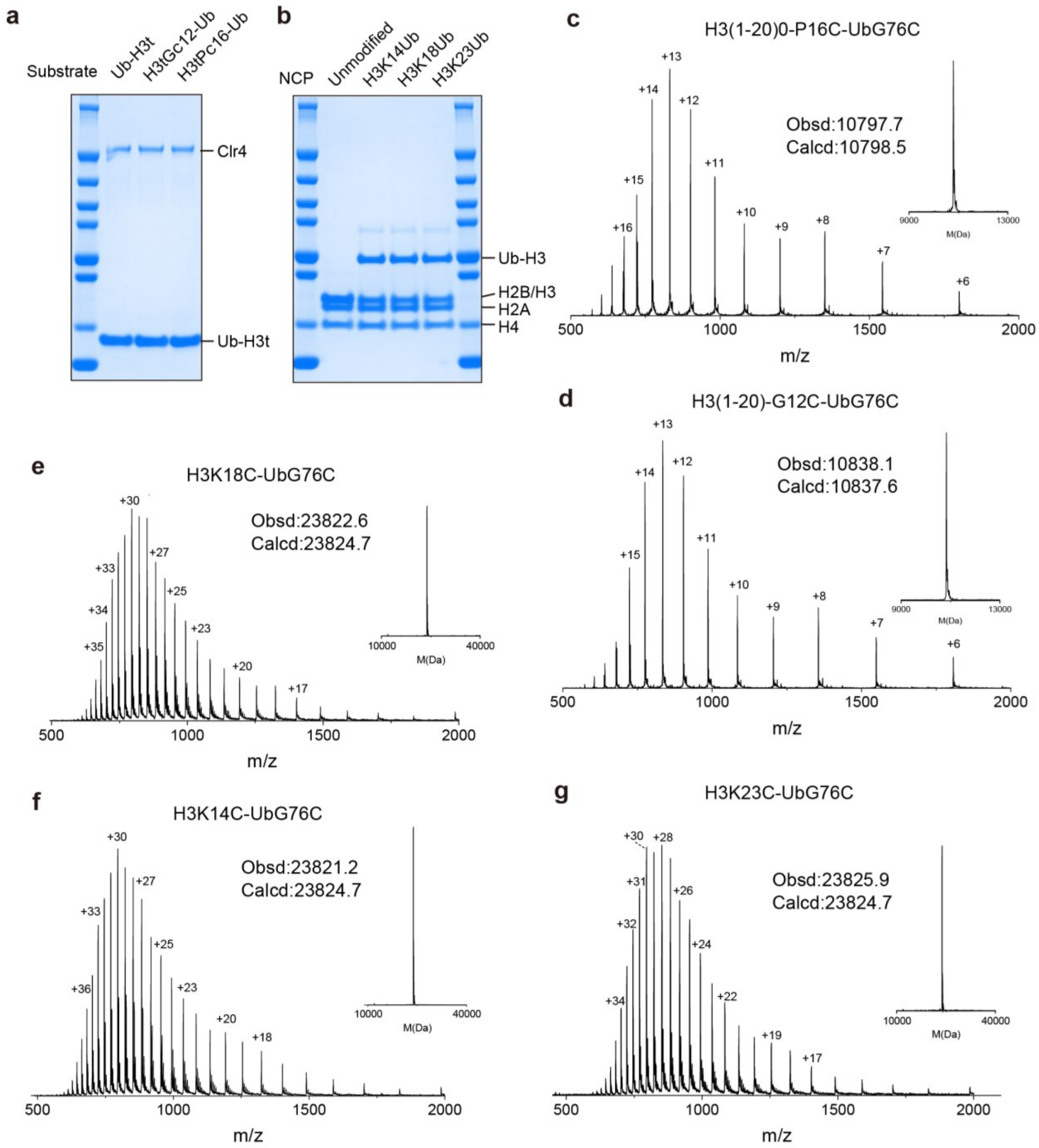


**Supplementary Fig. 8 | Property characterization of substrate samples for exploration of H3K14-selectivity for ubiquitination-mediated stimulation. a.** SDS-PAGE of the reaction system of WT Clr4 variants catalyzing Ub-H3t, H3tG$_{C}$12-Ub, and H3tP$_{C}$16-Ub methylation. **b.** SDS-PAGE of the unmodified NCP, NCP$_{H3K14Ub}$, NCP$_{H3K18Ub}$, and NCP$_{H3K23Ub}$. **c, d.** Representative ESI-MS traces of **c)** H3tG$_{C}$12-Ub, **d)** H3tP$_{C}$16-Ub. **e-g.** Representative ESI-MS traces of **e)** H3$_{K14C}$-Ub$_{G76C}$, **f)** H3$_{K18C}$-Ub$_{G76C}$, **g)** H3$_{K23C}$-Ub$_{G76C}$.
